# Targeting PFKFB3 Metabolic Gatekeeping of β-Cell Dysfunctional States Enables Durable Glucose Tolerance in Type 2 Diabetes

**DOI:** 10.64898/2026.09.20.752954

**Authors:** Kavit Raval, Erwin Ilegems, Alexandra C. Title, Emme Fishburn, Gurprit Bhardwaj, Olivier Mirguet, Paul Ratcliffe, Oppel Greeff, Abraham Van Wyk, Slavica Tudzarova

## Abstract

Progressive β-cell dysfunction underlies type 2 diabetes (T2D), yet existing therapies fail to correct the dysfunctional cellular states that accumulate during disease progression. We show that pharmacological inhibition of PFKFB3 markedly reduced the population of PFKFB3⁺ dysfunctional β-cells while preserving functional islet mass, consistent with selective depletion or phenotypic reprogramming of compromised β-cells. In transgenic diabetic mice expressing human IAPP, PFKFB3 inhibitor AZ67 improved glucose-stimulated insulin secretion and glucose tolerance through a mechanism consistent with direct restoration of β-cell function and distinct from incretin-based therapy. Critically, glycemic improvement persisted after treatment withdrawal, consistent with durable remodeling of islet functional states rather than transient pharmacological suppression. In human islet microtissues under glucotoxic and cytokine stress, AZ67 and its analogue AZ26 improved insulin secretion and proinsulin processing in a stress- and compound-dependent manner while reducing PFKFB3⁺ dysfunctional endocrine populations. Transcriptomic profiling identified suppression of inflammatory programs and activation of cholesterol homeostasis and adaptive metabolic pathways. These findings establish PFKFB3 inhibition as a potential disease-modifying therapeutic strategy for T2D.

**One Sentence Summary:** Inhibiting PFKFB3 clears dysfunctional β-cells and durably restores glucose control in diabetic mice and human islets.

## INTRODUCTION

Despite substantial therapeutic advances, it remains difficult to maintain glycemic control in type 2 diabetes (T2D) (*1*). Oral hypoglycemic agents including metformin, sulfonylureas, thiazolidinediones, and SGLT2 inhibitors lower blood glucose by improving insulin sensitivity, stimulating insulin secretion (GSIS), or enhancing renal glucose excretion, but their effectiveness frequently declines as β-cell dysfunction progresses (*2*), necessitating treatment intensification or combination therapies (*3–7*). GLP-1 receptor agonists (GLP-1RA) provide more potent glucose-lowering benefits through multiple mechanisms including incretin-mediated insulin potentiation, glucagon suppression, and delayed gastric emptying (*8–13*). Their efficacy depends on continued administration and preserved β-cell function, with glycemic control and weight benefits rapidly reversing upon treatment discontinuation (*11, 14–18*). Emerging single-cell and genetic studies in T2D demonstrate that β-cell dysfunction reflects persistent heterogeneous cellular states arising from metabolic and transcriptional remodeling, highlighting a major unmet need for therapies that selectively target dysfunctional β-cell populations (*19–25*). Importantly, neither oral agents nor GLP-1 receptor agonists correct underlying β-cell pathology or reduce stress-adapted dysfunctional β-cell states that accumulate during disease progression (*8, 26–28*). In contrast, inhibiting PFKFB3 (6-phosphofructo-2-kinase/fructose-2,6-bisphosphatase 3) offers a mechanistically distinct approach. In our previous report, PFKFB3 inhibitor AZ67 improved glycemic control by selectively eliminating dysfunctional β-cells identified by elevated PFKFB3 and MAIP1 expression through a mechanism consistent with cell-fitness competition (CFC) with PFKFB3 expression returning to baseline following resolution of cellular stress (*29*).

PFKFB3 is a bifunctional kinase/phosphatase regulating glycolytic flux, stress-driven metabolic reprogramming, and activity of inositol 1,4,5-trisphosphate receptor (IP3R) (*30–32*). Cross-phenotype GWAS analyses have implicated loci associated with glycolytic regulation, including regions linked to PFKFB3, in susceptibility to T1D, T2D, and latent autoimmune diabetes in adults (LADA) (*33*). Evidence has emerged that these effects are mediated through epistatic interactions across β-cell identity, inflammatory, and stress-response pathways, positioning metabolic control as a shared axis of genetic risk (*29, 33*). Consistent with this stress-associated phenotype, recent analysis of human pancreatic tissue demonstrated an increased proportion of PFKFB3⁺ β-cells in individuals with T2D, with PFKFB3 positivity inversely associated with β-cell mass and positively associated with islet amyloid deposition and pancreatic fibrosis (*34*). Under chronic glucotoxic and inflammatory stress, β-cells upregulate PFKFB3 as part of a survival glycolytic shift that supports ATP maintenance at the expense of stimulus-secretion coupling (*35, 36*), involving the calcium-regulating Matrix AAA Peptidase Interacting Protein 1 (MAIP1) (*29, 37*). These findings further support PFKFB3 as a marker of a pathologically stressed β-cell state in human diabetes. MAIP1 regulates mitochondrial Ca²⁺ homeostasis by facilitating assembly of the EMRE-containing mitochondrial calcium uniporter (MCU) complex, thereby controlling mitochondrial Ca²⁺ uptake and its dysregulation leads to calcium overload in mitochondria (*37*). Our epistasis analysis identified MAIP1 as functionally associated with PFKFB3 in dysfunctional β-cells, linking PFKFB3-dependent metabolic stress to mitochondrial Ca²⁺ regulation and providing a rationale for assessing MAIP1 as a complementary marker of the PFKFB3⁺ dysfunctional β-cell state (*29*). Genetic conditional knockdown of PFKFB3 selectively reset dysfunctional β-cells while sparing healthy β-cells consistent with activation of CFC pathways (*38*). Thus, PFKFB3 functions as a central metabolic node and a gatekeeper of CFC that protects dysfunctional β-cells from attrition while simultaneously sustaining their pathological, low-fitness phenotype and positing PFKFB3 as a mechanistic target for disease-modifying therapy (*29, 38*). Here, we tested whether pharmacological PFKFB3 inhibition produces durable restoration of β-cell function beyond active drug exposure and whether its mechanism is distinct from or complementary to GLP-1 receptor agonism. We characterized the biochemical and physicochemical properties of AZ67, compared its efficacy with Exendin-4 in hIAPP transgenic mice exposed to high-fat diet, and assessed durability following treatment withdrawal. We then evaluated PFKFB3 inhibition in human islet microtissues (hIsMT) exposed to glucotoxic or inflammatory stress and used transcriptomic profiling to define the stress-dependent molecular programs.

We show that AZ67 improves glucose tolerance and insulin secretion in vivo, consistent with selective depletion or phenotypic reprogramming of PFKFB3^+^ β-cells, and reversal of metabolic stress in hIsMT. Our results indicate that PFKFB3 inhibition remodels dysfunctional β-cell states and produces durable glycemic benefits that persist after treatment withdrawal, supporting its potential as a disease-modifying therapeutic approach.

## RESULTS

### PFKFB3 inhibition improves glucose tolerance and β-cell function through a mechanism distinct from incretin therapy

To evaluate pharmacological inhibition of PFKFB3 in vivo in the context of diabetes, we used human islet amyloid polypeptide (hIAPP) transgenic (h-TG) mice exposed to a high-fat diet (HFD), together with AZ67, a PFKFB3 inhibitor supported by extensive experimental evidence of on-target PFKFB3 engagement without adverse effects (*39–41*). Hemizygous expression of the hIAPP transgene in β-cells renders male h-TG mice susceptible to human-like diabetes upon HFD exposure, marked by progressive β-cell dysfunction, impaired first-phase insulin secretion, and glucose intolerance (*38*).

To establish translational suitability for in vivo studies, we first performed comprehensive biochemical and biophysical profiling of AZ67 and compared it with its analogue AZ26 (Fig. S1 and Table 1) (*42*). We determined LogD, solubility, plasma protein binding, plasma stability, and IC_50_ values of 10.5 nM and 29.6 nM for AZ67 and AZ26, respectively, consistent with previous reports (Fig. S1A-C) (*42*). Direct biophysical interaction of AZ67 and AZ26 with PFKFB3 was demonstrated using spectral-shift binding assays (SpS) (*43*). SpS measures ligand-induced changes in the intrinsic fluorescence of the target protein to determine equilibrium dissociation constants (Kd) independently of enzymatic activity (Fig. S1A). Both compounds bound PFKFB3 with nanomolar affinity (Kd ∼52 nM for AZ67 and 79 nM for AZ26), confirming direct physical engagement with the target (Fig. S1A). Although AZ67 displayed approximately 1.5-fold higher binding affinity than AZ26, both values fell within the nanomolar range and were consistent with their respective enzymatic IC_50_ values (Fig. S1A-C). Thus, both compounds directly engage PFKFB3 with nanomolar affinity and inhibit its enzymatic activity with comparable potency. Potential off-target kinase effects were evaluated using a KinaseProfiler panel comprising 390 kinases (Fig. S1D, E). Both AZ67 and AZ26 exhibited limited off-target kinase activity, with inhibition not exceeding 30% across the tested kinase panel (Fig. S1D,E). The top 20 kinases demonstrating the highest degree of inhibition by AZ67 and AZ26 are presented in Fig. S1D, E.

**Table 1.** Protein binding of AZ67 and AZ26 in human islet microtissue culture medium. The table represents the free fraction available, IC_50_ fold coverage and IC_90_ fold coverage at 5 µM concentration of AZ67 and AZ26.

|  |  |  | %Protein Binding |  |  | % Recovery |  |  | % Free fraction | Free fraction (μM) | Fold IC50 coverage | Fold IC90 coverage |
| --- | --- | --- | --- | --- | --- | --- | --- | --- | --- | --- | --- | --- |
| Compound ID | Concentration (μM) | Matrix | Replicate 1 | Replicate 2 | Mean | Replicate 1 | Replicate 2 | Mean |  |  |  |  |
| AZ 67 | 5 | Culture Medium | 83.6 | 86.7 | 85.2 | 87.1 | 100.5 | 93.8 | 14.8 | 0.74 | 70.4 | 7.83 |
| AZ 26 | 5 | Culture Medium | 86.9 | 89.0 | 87.9 | 104.5 | 92.1 | 98.3 | 12.1 | 0.60 | 20.3 | 2.25 |

Functional suppression of PFKFB3 by AZ67 was further demonstrated by a concentration-dependent reduction in fructose-1,6-bisphosphate (F1,6BP) across the higher AZ67 concentrations in MiaPaCa pancreatic cancer cells treated with 5-100 μM AZ67 using ion chromatography–mass spectrometry (IC-MS) (Fig. S2A).

Given our previous study demonstrating β-cell reset upon conditional β-cell specific PFKFB3 knockdown (*38*), we next investigated whether PFKFB3 inhibition engages a mechanism complementary to GLP-1 receptor agonism (GLP-1RA). Comparator studies were conducted in h-TG+HFD mice, (referred to hereafter as diabetogenic stress, DS) (Fig. 1). Exendin-4 (Ex-4), a well-characterized GLP-1RA widely used as an experimental benchmark for incretin-based therapy (*44*), served as the comparator over a 4-week treatment period, following the experimental design outlined in Fig. 1A.

**Fig. 1.**
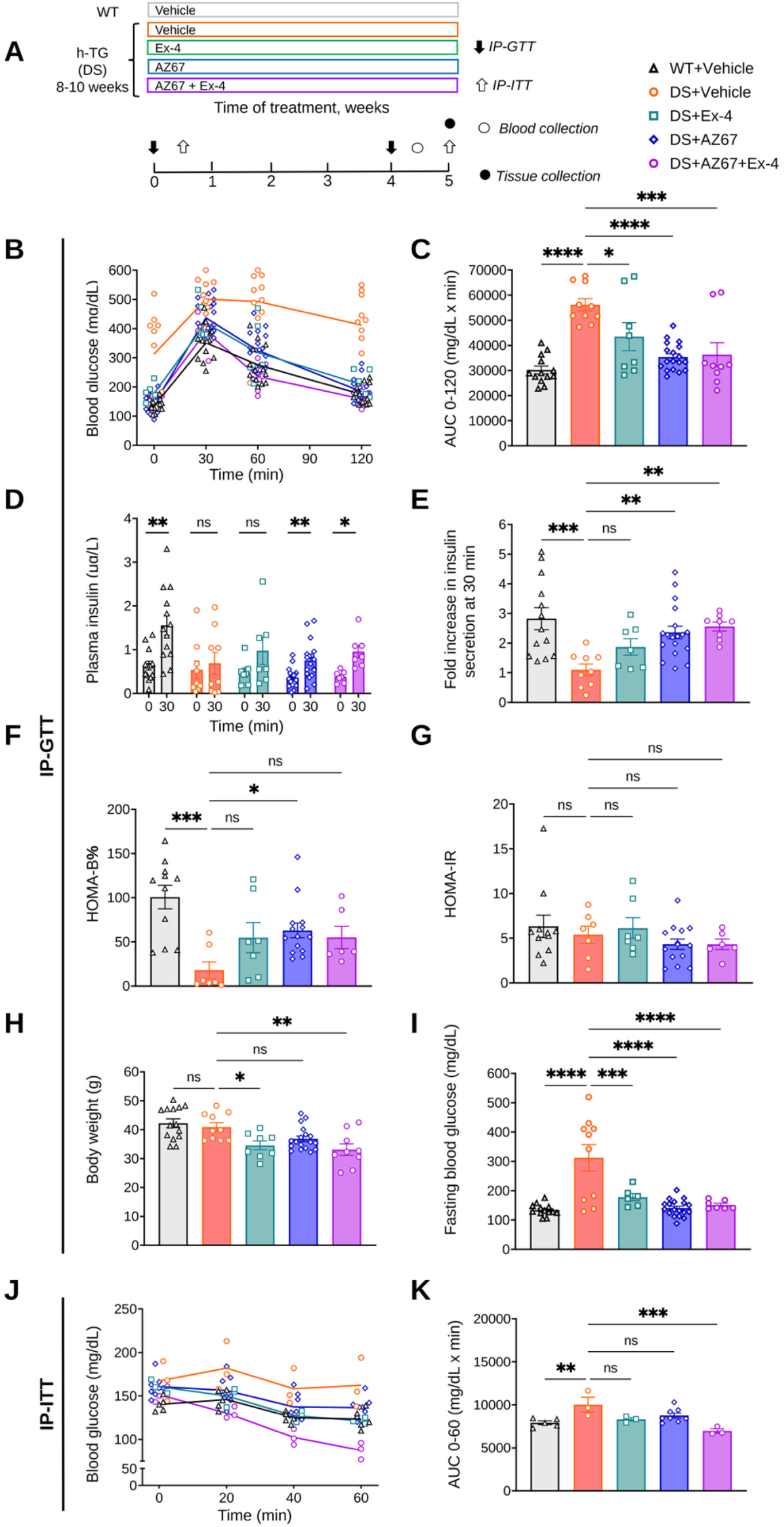
AZ67 restores glucose homeostasis and in vivo glucose-stimulated insulin secretion with additive effects in combination with Exendin-4. **(A)** Experimental design for in vivo comparator studies. All mice were fed High Fat Diet (HFD) and were assigned to the treatment groups at 8–10 weeks of age. Wild-type (WT) mice served as obese non-diabetic controls (WT+vehicle; n = 13–14). Human IAPP transgenic (h-TG) mice under HFD-induced diabetogenic stress (DS) were randomized to be injected with vehicle (DS+vehicle; n = 8–12) or treated with Exendin-4 (20 μg/kg/day i.p.; DS+Ex-4; n = 7–8), AZ67 (28 mg/kg/day i.p.; DS+AZ67; n = 18), or the combination (DS+AZ67+Ex-4; n = 7–9). AZ67 and Ex-4 were administered at least 6 hours apart. After 4 weeks, half of the AZ67 cohort was taken off treatment (DS+AZ67_G1; n = 9) or the rest received an additional 4 weeks of daily treatment (DS+AZ67_G2; n = 9) to assess durability. **(B)** Intraperitoneal glucose tolerance test (IP-GTT) glucose excursion curves following 4 weeks of treatment. **(C)** Area under the curve (AUC_0–120_) analysis for IP-GTT. **(D)** Plasma insulin concentrations at baseline and 30 minutes following intraperitoneal glucose challenge. **(E)** Fold increase in insulin secretion at 30 minutes following glucose challenge across treatment groups. **(F)** Homeostasis Model Assessment of β-Cell Function (HOMA-B%) calculated from fasting plasma insulin and blood glucose levels from IP-GTT after 4 weeks of AZ67 treatment. **(G)** Homeostasis Model Assessment of Insulin Resistance (HOMA-IR) calculated from fasting plasma insulin and blood glucose levels from IP-GTT after 4 weeks of AZ67 treatment. **(H)** Body weight (g) measurements across treatment groups after 4 weeks, as described in (A). **(I)** Fasting blood glucose (mg/dL) measured prior to IP-GTT after 4 weeks of treatment. **(J)** Intraperitoneal insulin tolerance test (IP-ITT) glucose excursion curves following 4 weeks of treatment. **(K)** Area under the curve (AUC_0–60_) analysis for IP-ITT. Statistical significance was determined using one-way ANOVA followed by Dunnett’s multiple-comparisons test. Non-significant comparisons are indicated as ns. Statistical significance is denoted as follows: P < 0.05 (*), P < 0.01 (**), P < 0.001 (***), and P < 0.0001 (****).

AZ67 significantly reduced IP-GTT AUC relative to DS+Vehicle (****P < 0.0001), with Ex-4 producing a smaller but significant improvement (*P < 0.05) and combination treatment also improving glucose tolerance (***P < 0.001; Fig. 1B,C). AZ67 significantly increased glucose-stimulated plasma insulin relative to baseline (**P < 0.01, Fig. 1D). Fold insulin stimulation was increased by AZ67 and AZ67+Ex-4 combination (**P < 0.01, Fig. 1E). AZ67 monotherapy significantly improved HOMA-β% relative to DS+Vehicle without altering HOMA-IR (*P < 0.05) (Fig. 1F,G). At four weeks, Ex-4, alone or in combination with AZ67, significantly reduced body weight and IP-ITT AUC was significantly reduced by AZ67+Ex-4 combination (***P < 0.001, Fig. 1J,K). Notably, the increase in HOMA-β% with AZ67 monotherapy, in the absence of a change in HOMA-IR, indicated that the metabolic improvement was primarily associated with enhanced β-cell function rather than altered peripheral insulin sensitivity.

Given that PFKFB3 genetic depletion (*38*) as well as inhibition by AZ67 (*29*) engage mechanisms consistent with cell fitness competition and are associated with depletion of the PFKFB3⁺ β-cell population concomitant with recovery of functional β-cell representation, we reasoned that this unique mechanism might confer efficacy that persists beyond the active drug exposure.

### PFKFB3 inhibition produces durable glycemic improvement beyond active drug exposure

To determine whether PFKFB3 inhibition provides efficacy beyond active drug exposure and thus therapeutic differentiation relative to Ex-4, we tested its effect on glucose tolerance after therapy withdrawal. We compared continuous treatment of AZ67 (G2, 8 weeks) versus an on-off regimen (G1, 4 weeks treatment followed by 4 weeks withdrawal) in h-TG+HFD (DS) mice (Fig. 2A-G). Continuous AZ67 treatment for 8 weeks restored glucose tolerance, reducing IP-GTT AUC relative to stress DS+Vehicle controls (****P < 0.0001; Fig. 2B,C). Notably, mice treated with AZ67 for 4 weeks followed by 4 weeks off-treatment retained comparable glycemic improvement at the study endpoint (****P < 0.0001 vs DS+Vehicle), demonstrating persistence of the glycemic benefit beyond active drug exposure. Ex-4 monotherapy for 8 weeks restored glucose tolerance and insulin sensitivity compared to DS+Vehicle (Fig. 2B-G). At 8 weeks, continuous AZ67, Ex-4, and AZ67+Ex-4 treatment each improved glucose tolerance relative to DS+Vehicle, with the combination producing the largest reduction in IP-GTT AUC (Fig. 2B,C). AZ67 (on-off and continuous) reduced fasting glucose levels (**P < 0.01 both; Fig. 2E) without affecting body weight (ns both; Fig. 2D), consistent with a mechanism primarily associated with improved β-cell function rather than systemic metabolic compensation.

**Fig. 2.**
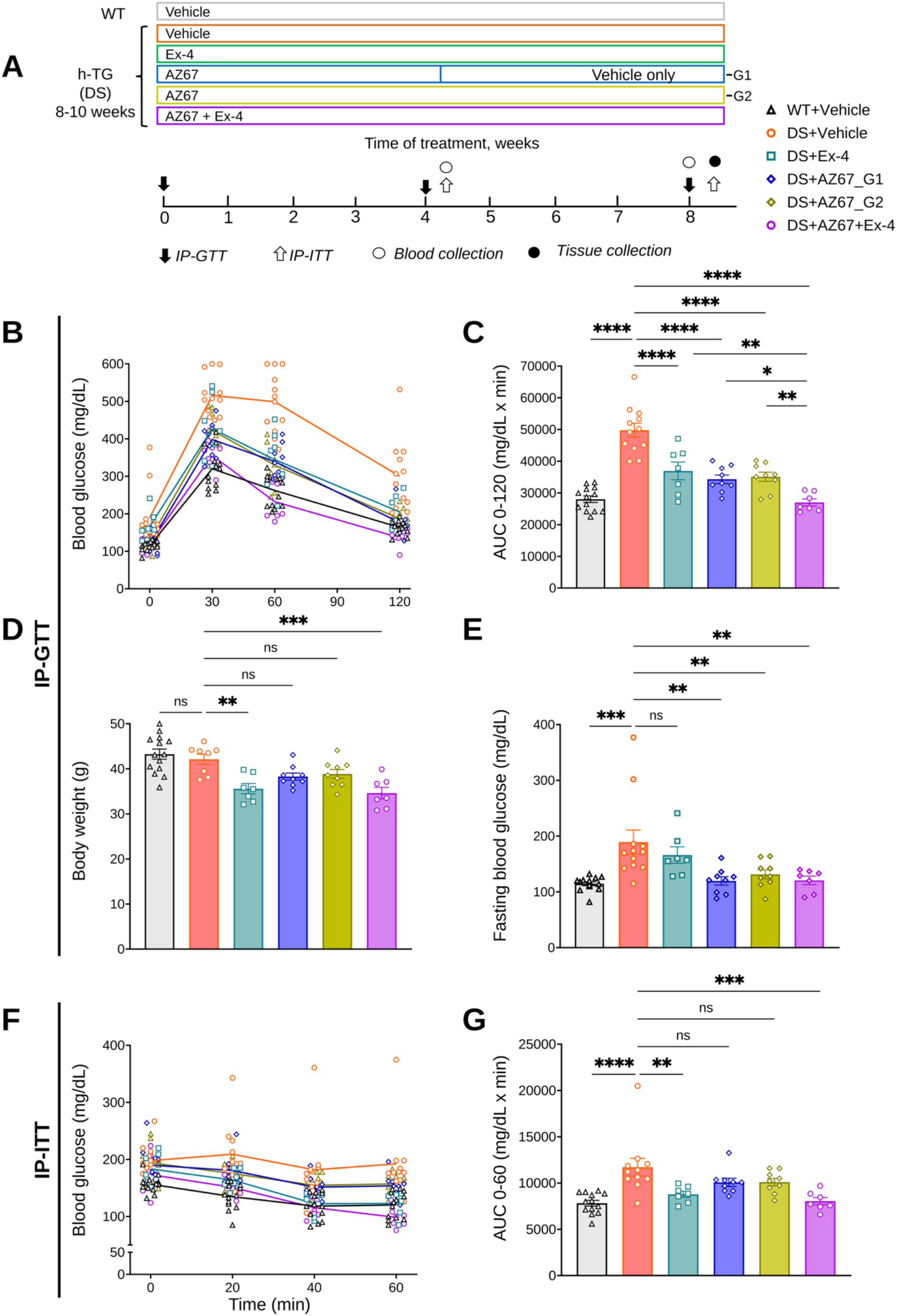
AZ67 induces durable glycemic control following treatment withdrawal in h-TG mice. **(A)** Following the 4-week comparative efficacy analysis described in Fig. 1A, AZ67-treated h-TG+HFD mice were subdivided into two cohorts to assess treatment durability: an “on-off” cohort receiving AZ67 for 4 weeks followed by 4 weeks without treatment (DS+AZ67_G1) and a continuous-treatment cohort receiving AZ67 for 8 weeks (DS+AZ67_G2). Exendin-4 (Ex-4) monotherapy and combination treatment groups continued their respective regimens for the full 8-week study period for comparative analyses. **(B)** Intraperitoneal glucose tolerance test (IP-GTT) glucose excursion curves following 8 weeks of treatment. **(C)** Area under the curve (AUC₀–₁₂₀) analysis for IP-GTT. **(D)** Body weight (g) measurements across treatment groups after 8 weeks. **(E)** Fasting blood glucose measurements across treatment groups after 8 weeks. **(F)** Intraperitoneal insulin tolerance test (IP-ITT) glucose excursion curves following 8 weeks of treatment. **(G)** Area under the curve (AUC₀–₆₀) analysis for IP-ITT. Statistical significance was determined using one-way ANOVA followed by Dunnett’s multiple-comparisons test. Non-significant comparisons are indicated as ns. Statistical significance is denoted as follows: P < 0.05 (*), P < 0.01 (**), P < 0.001 (***), and P < 0.0001 (****).

We next asked whether the cellular reset in PFKFB3^+^ and MAIP1^+^ β-cell frequency (*37*) may explain the differences between AZ67 and Ex-4.

### AZ67 selectively reduces PFKFB3^+^ β-cell population in vivo

β-cells upregulate PFKFB3 as part of a survival glycolytic shift, and its co-expression with MAIP1 in dysfunctional β-cells implicates both proteins in the maladaptive metabolic state (*29*). To determine whether the observed durable efficacy is a reflection of remodeled β-cell phenotype, pancreatic tissue sections were analyzed by immunofluorescence staining for PFKFB3 and MAIP1 (Fig. 3A-D) (split images shown in Fig. S3 and S4). DS+Vehicle mice exhibited an increased population of PFKFB3^+^ β-cells (47.03%, ****P < 0.0001, n=3), consistent with the persistent metabolically compromised β-cells observed in human diabetes (*35*). Both continuous (G2) and on-off (G1) AZ67 treatment regimens reduced the fraction of PFKFB3^+^ β-cells to less than 1% (****P < 0.0001), accompanied by recovery of the β-cell proportion within islets. The number of β-cells relative to total islet cells in DS+Vehicle mice was reduced relative to WT controls, consistent with progressive β-cell loss under DS (Fig. 3E). On-off AZ67 monotherapy (DS+AZ67_G1), continuous AZ67 regimen (DS+AZ67_G2) and the AZ67+Ex-4 combination each achieved significant recovery of β-cell proportion relative to total islet cells (Fig. 3E). Ex-4 monotherapy did not significantly expand β-cell proportion relative to DS+Vehicle (Fig. 3E). This was consistent with the incomplete restoration of in vivo GSIS at 4 weeks by Ex-4 monotherapy (ns; Fig. 1D, E). Combination therapy resulted in a greater reduction in PFKFB3^+^ β-cells (****P < 0.0001), indicating complementary mechanisms (Fig. 3A, C). MAIP1⁺ β-cell population, associated with β-cell calcium toxicity (*37*), was reduced across all treatment groups including Ex-4 monotherapy (Fig. 3B, D).

**Fig. 3.**
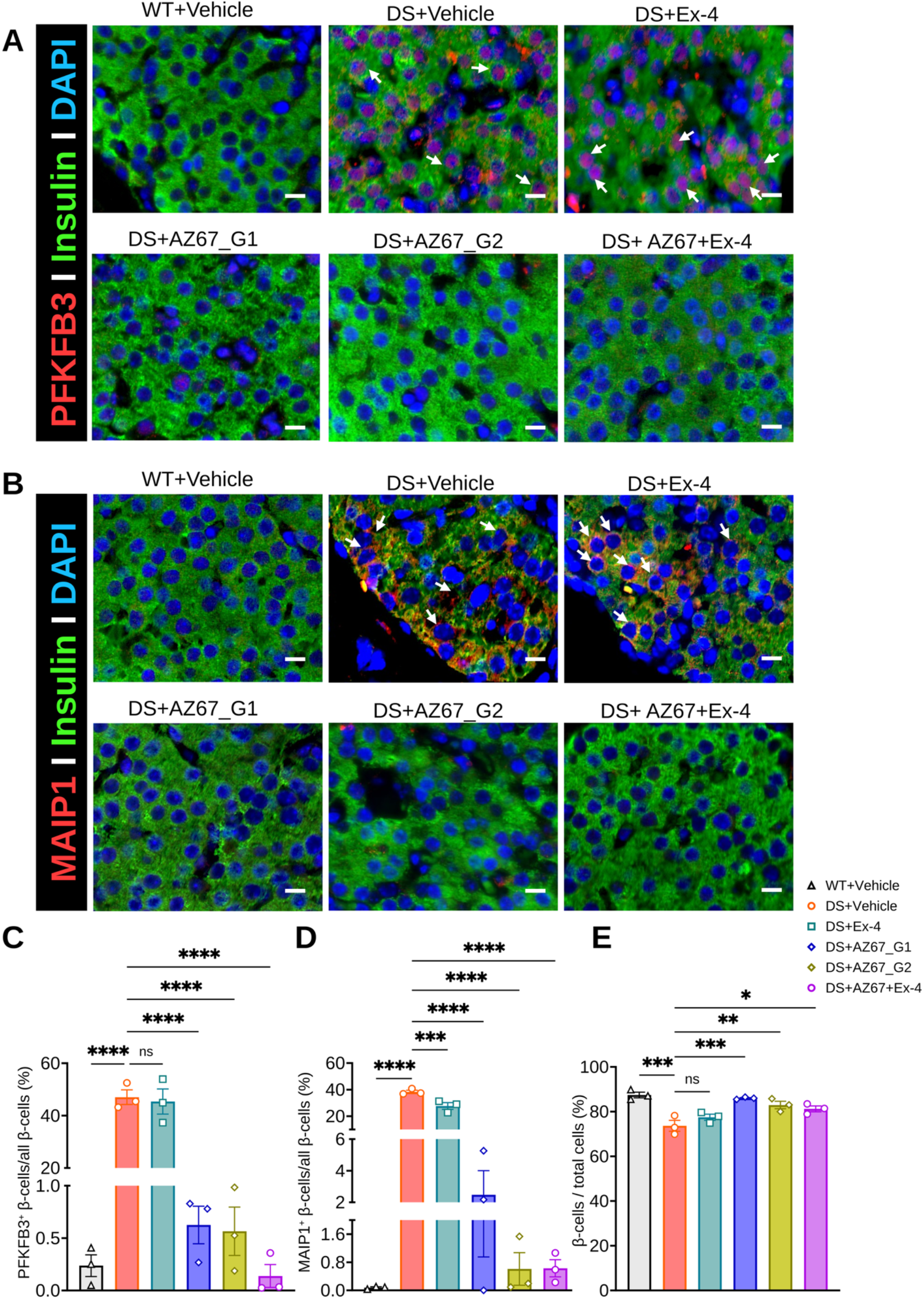
AZ67 reduces PFKFB3⁺ and MAIP1⁺ dysfunctional β-cell populations in diabetic h-TG mice. **(A)** Representative immunofluorescence images of pancreatic sections stained for PFKFB3 (red), insulin (green), and DAPI (blue) across treatment groups. AZ67 treatment under both continuous and on-off regimens reduced the abundance of PFKFB3⁺ dysfunctional β-cells, whereas Exendin-4 monotherapy had minimal effect (see inset). The greatest reduction was observed with combination therapy (AZ67 + Ex-4). Scale bar: 10 μm. **(B)** Representative immunofluorescence images of pancreatic sections stained for MAIP1 (red), insulin (green), and DAPI (blue) across treatment groups. AZ67 (continuous and on-off regimens), Ex-4 monotherapy, and combination treatment all reduced the abundance of MAIP1⁺ dysfunctional β-cells. **(C)** Quantification of PFKFB3^+^ β-cells as a percentage of total INS^+^ β-cells across treatment groups. **(D)** Quantification of MAIP1^+^ β-cells as a percentage of total INS^+^ β-cells across treatment groups. **(E)** Quantification of INS^+^ β-cells as a percentage of total DAPI^+^ islet cells across treatment groups. Data are presented as mean ± SEM (n = 3 mice per group; 25–40 islets analyzed per pancreas). Statistical analysis was performed using one-way ANOVA followed by Dunnett’s post hoc multiple-comparisons test versus DS+Vehicle. Statistical significance is indicated as follows: P < 0.05 (*), P < 0.01 (**), P < 0.001 (***), and P < 0.0001 (****).

We next sought to understand whether the effect of PFKFB3 inhibition is reproduced in human islet hIsMT exposed to T2D- and T1D-like stress conditions.

### Human islet microtissues (hIsMT) confirm translational relevance of PFKFB3 inhibition across glucotoxic and inflammatory stress models

To evaluate human translational relevance, AZ67 and AZ26 were tested in 3D human islet hIsMT (human microtissues, hIsMT) composed of primary endocrine cells (Fig. 4) (*45–47*). We used IC_50_ as well as protein binding and free fraction properties of the PFKFB3 inhibitors (AZ67 and AZ26) in the culture media for hIsMT to calculate treatment concentration in vitro (Fig. S1 and Table 1). hIsMT were chronically exposed to glucotoxic (GTX, 11 mM glucose for 6 days) or cytokine-induced pro-inflammatory stress (CYTO, 2 ng/mL IL-1β, 10 ng/mL TNFα, 10 ng/mL IFNγ) to model ER- and inflammatory disease components common to T2D and T1D (*48, 49*) (Fig. 4). Both stress culture conditions induced hallmark β-cell dysfunction, including impaired GSIS, reduced insulin content, increased basal insulin secretion and proinsulin-to-insulin ratios, and expansion of PFKFB3⁺ endocrine cell populations (Figs. 4–6 and Figs. S5 and S6).

**Fig. 4.**
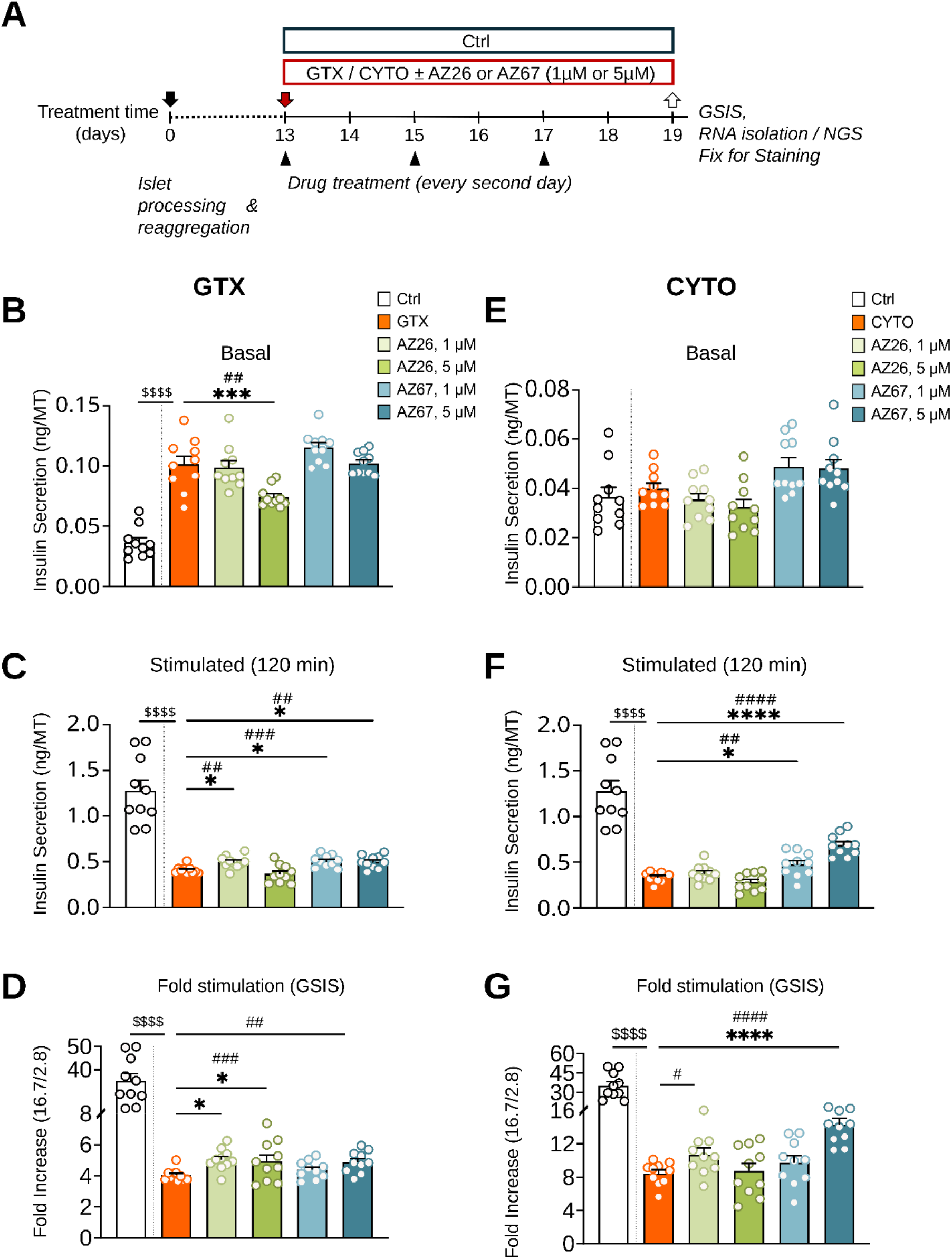
AZ67 and AZ26 modulate β-cell function and reduce metabolic stress in human islet hIsMT exposed to glucotoxic and cytokine stress. **(A)** Treatment protocol: human islet microtissues (hIsMTs) were cultured under glucotoxic conditions (11 mM glucose) or under cytokine-induced stress (IL-1β, TNFα, and IFNγ) for 6 days in the presence or absence of AZ67 or AZ26 (1 μM or 5 μM). **(B)** and **(E)** basal insulin secretion in GTX and CYTO, respectively. **(C)** and **(F)** (GSIS) secretion per MT following 120-min glucose stimulation in GTX and CYTO, respectively. **(D)** and **(G)** fold stimulation GSIS following 16.7 mM glucose in GTX and CYTO, respectively. Data are presented as mean ± SEM (n = 8–10 independent microtissues). Ratios were calculated using molar concentrations. Following outlier removal using ROUT analysis (Q = 5%), one-way ANOVA followed by Dunnett’s multiple-comparisons test was used for prespecified comparisons of each treatment group with the corresponding GTX or CYTO stress control (stars). Exploratory pairwise comparisons were additionally performed using unpaired two-tailed t-tests (hash symbols). Unpaired t-tests comparing healthy control microtissues cultured under standard glucose conditions (2.8 mM) versus glucotoxic conditions were performed to validate disease induction (dollar symbols). P < 0.05 (*/#/$), P < 0.01 (**/##/$$), P < 0.001 (***/###/$$$), and P < 0.0001 (****/####/$$$$).

**Fig. 5.**
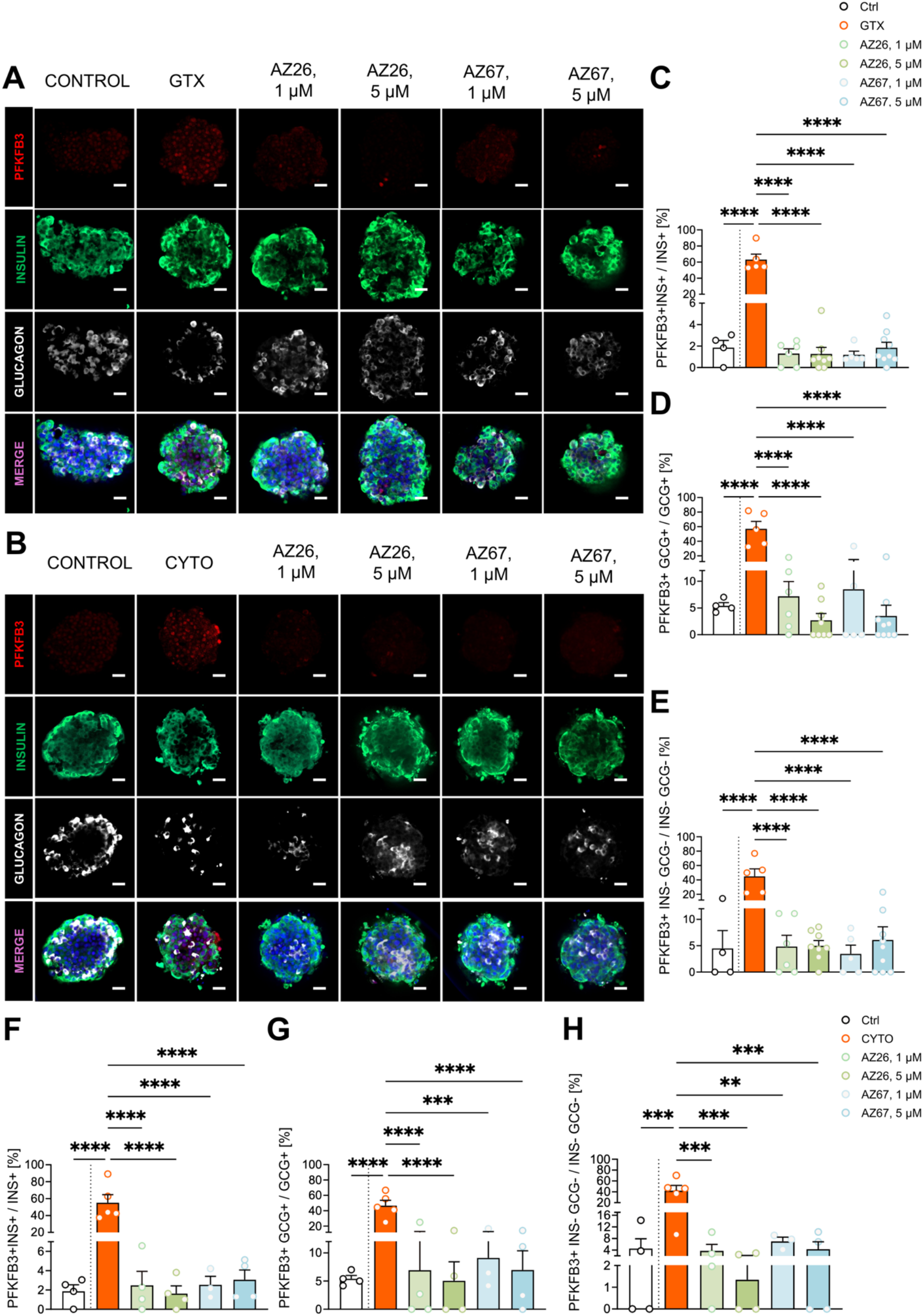
Effects of AZ67 and AZ26 on PFKFB3⁺ endocrine cell populations in human islet hIsMT under diabetogenic stress. Representative immunofluorescence images of hIsMT stained for insulin (green), glucagon (white), PFKFB3 (red), and nuclei (blue) in **(A)** glucotoxicity-induced (GTX) and **(B)** cytokine-induced (CYTO) models of diabetes, demonstrating reduced PFKFB3⁺ cell populations following treatment with AZ67 and AZ26. A representative middle optical z-section from the hIsMT z-stack is shown, selected to capture the maximal cross-sectional area of the hIsMT. Scale bar: 20 μm. **(C)** and **(F)** PFKFB3⁺ INS⁺ β-cells relative to total INS⁺ β-cells in GTX or CYTO, respectively. **(D)** and **(G)** PFKFB3⁺ GCG^+^ α-cells relative to total GCG^+^α-cells in GTX or CYTO, respectively. **(E)** and **(H)** PFKFB3⁺ INS⁻GCG⁻ double-negative endocrine cells relative to total INS⁻GCG⁻ double-negative cells in GTX or CYTO, respectively. Quantification was performed using QuPath software. Quantification represents the mean of three optical z-sections sampled from the top, middle, and bottom regions of each hIsMT. Data are presented as mean ± SEM (n = 4–9 independent microtissues for GTX and n = 3–5 independent microtissues for CYTO). Statistical significance was determined using one-way ANOVA followed by Dunnett’s multiple-comparisons test. Statistical significance is indicated as follows: P < 0.05 (*), P < 0.01 (**), P < 0.001 (***), and P < 0.0001 (****).

**Fig. 6.**
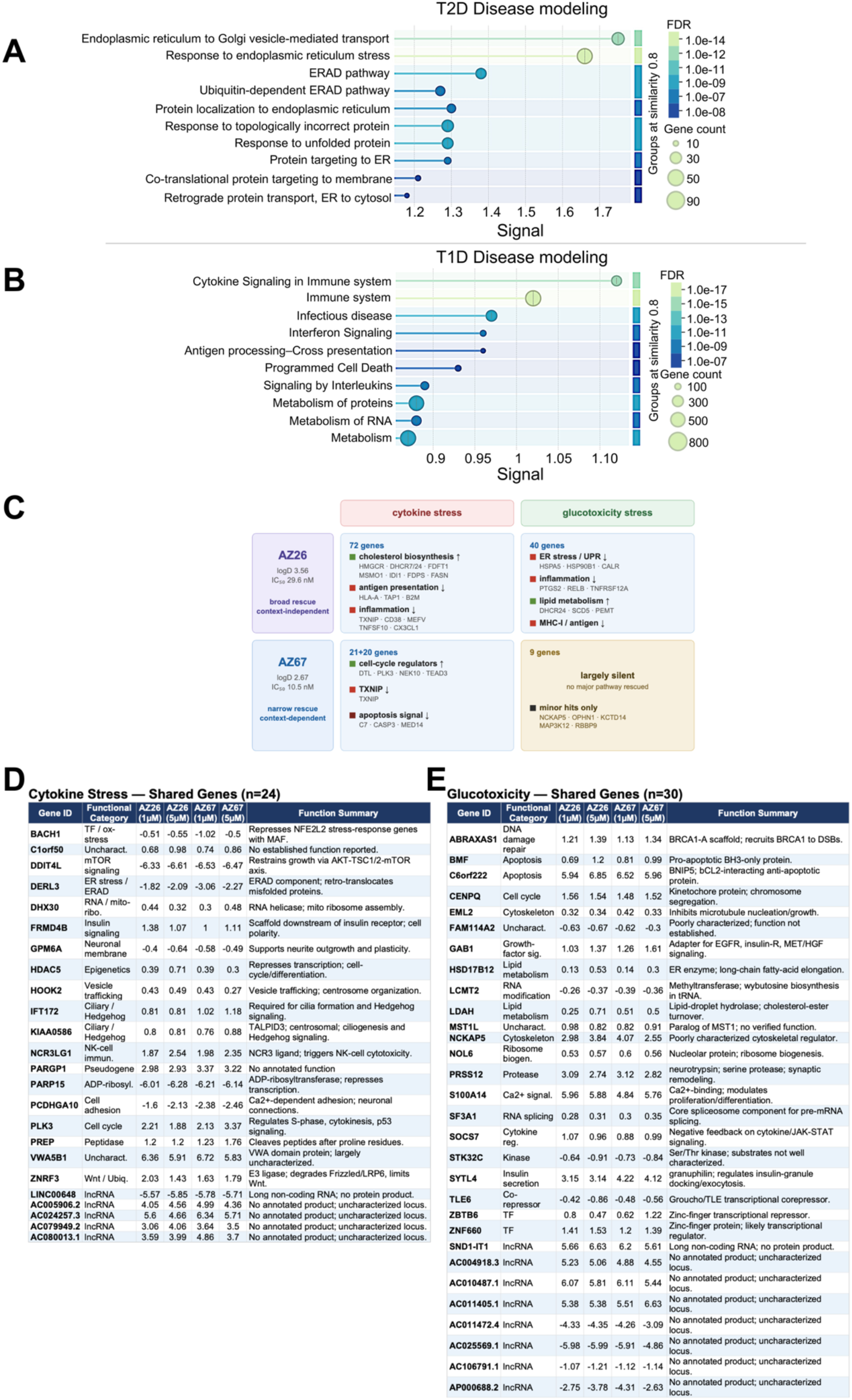
Transcriptional rescue signatures of AZ26 and AZ67 in stressed human islet hIsMT. **(A)** GSEA of DEGs under GTX stress showing enrichment of ER stress, unfolded protein response, and ubiquitin-dependent degradation pathways. Bubble size, gene count; color, FDR. **(B)** GSEA under CYTO stress showing enrichment of cytokine and interferon signaling, antigen presentation, and programmed cell death pathways, confirming disease-relevant modeling. **(C)** Summary of reversed genes across GTX and CYTO contexts. **(D)** Functional summary of the 24 genes identified as overlapping differentially expressed genes (DEGs) between AZ26 and AZ67, independently at both 1 µM and 5 µM, under cytokine stress (nominal unadjusted p < 0.05). For each gene, log2FC values are shown for both compounds at both concentrations, alongside a functional category and brief description (paraphrased from STRING/UniProt). **(E)** As in **(D)**, for the 30 overlapping DEGs identified between AZ26 and AZ67 under glucotoxicity (n=30; nominal unadjusted p <0.05).

PFKFB3 inhibition improved GSIS in a stress- and compound-dependent manner (Fig. 4). Under GTX stress, AZ26 significantly increased stimulation index by Dunnett’s multiple-comparisons analysis, whereas AZ67 showed evidence of improvement in pairwise analyses relative to GTX stress. Under CYTO stress, AZ67 produced the strongest functional rescue, with 5 μM AZ67 significantly increasing both stimulated insulin secretion and stimulation index (Fig. 4F,G). Under CYTO stress, both AZ26 and AZ67 at 5 μM significantly reduced the elevated basal proinsulin-to-insulin ratio, consistent with partial restoration of proinsulin processing (Fig. S6D); this effect was not reproduced consistently under GTX stress (Fig. S5D). AZ26 produced partial rescue across GSIS endpoints and was less effective than AZ67 under CYTO conditions, consistent with its approximately two-fold lower enzymatic potency (Fig. S1).

Immunofluorescence analysis demonstrated that AZ67 and AZ26 significantly reduced the PFKFB3⁺ fraction of INS⁺ β-cells, GCG⁺ α-cells, and INS⁻/GCG⁻ stressed endocrine populations in both GTX- and CYTO-treated hIsMT, consistent with the findings in pancreata of h-TG+HFD mice treated with AZ67 (Fig. 5A–H and Fig. 3C). Representative images of the middle optical z-section of the hIsMT are shown in Fig. 5A–B, while representative top and bottom optical z-sections from the corresponding z-stacks are shown in Figs. S8 and S9, respectively, in addition to a supporting video file for representative hIsMT per treatment group.

### PFKFB3 inhibition induces transcriptional programs associated with metabolic rewiring and immune quiescence

To define the transcriptional response to PFKFB3 inhibition in human islet hIsMT, we performed RNA-seq under glucotoxic (GTX) and cytokine (CYTO) stress with or without AZ67 or AZ26 (1 or 5 µM; Fig. 6). Read quality was confirmed (Fig. S7). STRING analysis showed that GTX preferentially induced ER-stress and proteostasis pathways associated with T2D-like β-cell dysfunction, whereas CYTO activated inflammatory and immune programs characteristic of T1D-associated injury (Fig. 6A,B; tables S1 and S2), validating both models as disease-relevant (*50, 51*).

CYTO produced a broader transcriptional response than GTX. Its signature was dominated by interferon/NF-κB-associated inflammation, including *IFI44L, CD38, MEFV*, and *TXNIP*, together with suppression of lipid metabolism, whereas GTX preferentially activated ER-stress genes (*HSPA5, HSP90B1*), *COX-2/PTGS2*, and MHC-I antigen-presentation genes (*NLRC5, HLA-B/E, TAP1*; Fig. 6A,B; tables S1 and S2).

Both inhibitors induced transcriptional reprogramming, with more DEGs at 5 µM. AZ26 produced a broader transcriptional response, reversing 72 genes under CYTO and 40 genes under GTX stress (Fig. 6C and Fig. S10). Across both conditions, AZ26 restored the SREBP2-dependent cholesterol/mevalonate program, including enzymes from *HMGCR* to *DHCR7*, defining a stress-independent lipid-homeostasis signature (Fig. 6C; tables S11 to S20). This is consistent with evidence that suppression of the SREBP2/cholesterol axis is an early feature of β-cell proteotoxicity in T2D (*52*). AZ26 also suppressed context-specific inflammatory programs, including *CD38* and *TNFSF10* under CYTO stress and the RELB–PTGS2–MHC-I axis under GTX stress.

AZ67 produced a more focused response. Under CYTO stress, both doses suppressed *TXNIP*, a mediator of oxidative stress and β-cell apoptosis (*53, 54*), whereas 1 µM AZ67 restored cell-cycle regulators including *DTL*, *PLK3*, *NEK10*, and *TEAD3*. Under GTX stress, AZ67 produced fewer transcriptional changes, indicating stronger context-dependent activity.

To identify compound-convergent, dose-robust responses, we compared DEGs nominally significant at unadjusted P < 0.05, that changed concordantly at 1 and 5 µM for both compounds. This identified 24 shared DEGs under CYTO and 30 under GTX stress (Fig. 6D,E). The two sets were largely functionally distinct. Under GTX, shared genes included the apoptosis regulators *BNIP5* (*C6orf222*) and *BMF*, together with the β-cell-relevant genes *SYTL4*, involved in insulin-granule docking and exocytosis (*55*), and *GAB1*, an adaptor in insulin- and growth-factor signaling (*56*). Under CYTO stress, convergence centered on *FRMD4B*, a signaling scaffold linked to insulin-receptor pathways (*57*). Because these candidate genes were defined using a nominal unadjusted significance threshold, these associations should be considered hypothesis-generating pending orthogonal validation or multiple-testing-corrected analysis.

Together, AZ26 induced broad metabolic and lipid-homeostasis remodeling across stress conditions, whereas AZ67 produced a more focused, stress-context-dependent response.

## DISCUSSION

The studies presented here converge on a mechanistic framework in which pharmacological inhibition of PFKFB3 alleviates metabolic stress, normalizes insulin secretory dynamics, and reshapes the composition of the β-cell population through mechanisms consistent with either β-cell fitness competition or downregulation of PFKFB3 protein levels. Rather than acting as a classical glucose-lowering intervention, PFKFB3 inhibition engages a biological program that targets the dysfunctional β-cell state itself, consistent with a disease-modifying therapeutic paradigm.

Dysfunctional β-cells in diabetes exhibit activation of stress-associated metabolic programs characterized by HIF1α activation, elevated PFKFB3 expression, calpain-mediated proteolysis including c-Myc truncation, MAIP1 expression, impaired proinsulin processing, and disrupted insulin exocytosis (*29, 35, 58–60*). These dysfunctional cellular states accumulate under proteotoxic, glucotoxic, lipotoxic, and inflammatory conditions and contribute to reduced first-phase insulin secretion and to β-cell dedifferentiation (*61–63*). Previous work demonstrated that PFKFB3 expression identifies a subpopulation of metabolically compromised β-cells exhibiting prolonged survival despite impaired secretory function (*29*). Importantly, independent analysis of human T2D pancreata has now shown increased PFKFB3⁺ β-cell abundance associated with reduced β-cell mass, amyloid deposition, and pancreatic fibrosis, further linking PFKFB3 expression to progressive pathological remodeling of the diabetic pancreas (*34*). Pharmacological inhibition of PFKFB3 appears to dismantle this maladaptive survival program similar to inhibition of its upstream regulator HIF1α (*64*), facilitating clearance or reprogramming of metabolically compromised β-cells.

Across in vivo studies, AZ67 consistently improved glucose tolerance and GSIS in h-TG+HFD diabetic mice. The magnitude of glycemic improvement, particularly in GSIS, was comparable to or exceeded that achieved with Ex-4, yet occurred without concomitant reduction in body weight or insulin sensitivity. AZ67 significantly rescued the diabetes-associated reduction in β-cell function (HOMA-B%, Fig. 1F) seen in DS+Vehicle mice, whereas HOMA-IR (Fig. 1G) remained unchanged across all groups, indicating that the DS phenotype and its rescue by AZ67 are driven primarily by a β-cell secretory defect rather than by peripheral insulin resistance. The dissociation from changes in body weight and insulin sensitivity supports a predominantly β-cell-centered mechanism of action. Restoration of in vivo GSIS is particularly notable because impairment of this response represents an early and persistent defect in diabetes pathophysiology that is insufficiently addressed by many currently available therapies (*5, 62, 65*).

Diabetic mice receiving AZ67 for four weeks retained significant metabolic improvement over four additional weeks without drug exposure, suggesting that PFKFB3 inhibition induces sustained remodeling of β-cell population composition, in contrast to Ex-4 treatment. Histological analyses demonstrated near-complete disappearance of PFKFB3^+^ β-cells following AZ67 treatment, reaching numbers similar to what was observed in WT β-cells. Although longer-term studies will be required to define the duration of this effect beyond the current observation window, the persistence of the glycemic correction following treatment discontinuation represents a key observation supporting durability and disease-modifying potential.

Human islet microtissue (hIsMT) studies recapitulated the central phenotypes observed in vivo, providing translational relevance. Under GTX and CYTO stress, human islet hIsMT exhibited reduced insulin secretion, impaired proinsulin processing, and expansion of PFKFB3^+^ endocrine populations. AZ26 and AZ67 improved basal proinsulin-to-insulin ratios under CYTO stress at 5 µM, suggesting partial restoration of proinsulin processing under inflammatory conditions; this effect was not consistently observed under glucotoxic stress, indicating stress context-dependent activity.

Although AZ26 and AZ67 share the same molecular target, PFKFB3, they produced distinct transcriptional profiles. AZ26 drove a broader response encompassing SREBP2-associated cholesterol and lipid biosynthesis genes across both stress contexts, alongside context-specific changes in immune and antigen presentation pathways (Fig. 6C, Fig. S10). AZ67 produced a narrower, context-dependent response, primarily affecting cell-cycle regulators and *TXNIP* under cytokine stress, with limited transcriptional changes under glucotoxic conditions (Fig. 6C), consistent with the restored β-cell numbers observed following AZ67 monotherapy in vivo. The selective rescue of cell cycle regulators (*DTL, PLK3, NEK10, TEAD3*) by AZ67 under cytokine stress raises the possibility that AZ67 may engage nuclear pools of PFKFB3, which has been reported to regulate cell cycle progression independently of its glycolytic function. This warrants further investigation with nuclear/cytoplasmic fractionation studies. The broader transcriptional reach of AZ26 may partly reflect its higher lipophilicity (logD7.4 3.56 vs. 2.67), potentially influencing intracellular accumulation or lipid membrane modification, which remains elusive. As profiling was performed on whole islet hIsMT, these signatures represent islet-wide responses, and cell type-resolved analyses will be needed to further interpret these findings. Compound-convergent DEG analysis (Fig. 6, D and E) showed that AZ26 and AZ67 converge on a stress-context-dependent core of PFKFB3-associated targets. Under glucotoxic stress, this included concurrent up-regulation of anti-apoptotic *BNIP5* and pro-apoptotic *BMF* consistent with remodeling of apoptosis-associated pathways in dysfunctional β-cells alongside the insulin-secretory genes *SYTL4* and *GAB1*. Under cytokine stress, convergence centered instead on the insulin-receptor scaffold *FRMD4B*.

Taken together, the integrated data support a mechanistically distinct paradigm in which PFKFB3 inhibition directly targets the dysfunctional β-cell state rather than compensating for downstream metabolic consequences, selectively reprogramming or eliminating compromised β-cells while preserving the secretory capacity of functional ones, thereby restoring coordinated insulin dynamics. This study has, however, limitations. The in vivo efficacy and durability studies were performed in a single male hIAPP-transgenic/HFD model, and durability was evaluated for four weeks after treatment withdrawal. Human islet organoid experiments were performed using microtissues derived from a single donor and therefore do not capture inter-donor variability. The reduction in PFKFB3⁺ β-cells is consistent with selective depletion or phenotypic reprogramming but does not distinguish these mechanisms directly.

All together, these findings support PFKFB3 inhibition as a promising potential disease-modifying therapeutic approach for diabetes and provide a rationale for further translational development.

## TRANSLATIONAL RELEVANCE

Progressive loss of functional β-cell capacity limits durable glycemic control in diabetes, whereas current therapies primarily compensate for rather than directly target dysfunctional β-cell states. Here, pharmacological PFKFB3 inhibition improved glucose-stimulated insulin secretion and glucose tolerance in a human IAPP transgenic model of diabetes and maintained glycemic benefit for four weeks after treatment withdrawal. PFKFB3 inhibition also reduced stress-associated PFKFB3⁺ endocrine populations and improved β-cell function in human islet microtissues exposed to glucotoxic or inflammatory stress. These findings support therapeutic targeting of PFKFB3-dependent metabolic stress as a strategy to remodel dysfunctional β-cell states and provide a rationale for developing PFKFB3 inhibitors as potential disease-modifying therapies for diabetes.

## MATERIALS AND METHODS

### Study Design

This study evaluated the PFKFB3 inhibitors AZ67 and AZ26 using a tiered experimental strategy encompassing in vitro ADME profiling, in vivo efficacy and durability studies, comparator analyses alone and in combination with Exendin-4, and translational studies in human islet microtissue modelling T1D and T2D-like diabetogenic stress. Animals were randomized to treatment groups as described below. Sample sizes, experimental units, statistical tests, and exclusion criteria are specified in the corresponding Methods sections and figure legends. Experiments were conducted in accordance with standard operating procedures at Eurofins Discovery Services for in vitro ADME profiling, InSphero for human islet microtissue studies, and UCLA Tudzarova Lab for efficacy, durability, and comparator analyses.

### Chemicals and Reagents

AZ67 (Cat. No. 5742) and its analogue AZ26 (Cat. No. 5675) were purchased from Tocris Biosciences. Exendin-4 (Cat. No. E7144) and 2-hydroxypropyl-β-cyclodextrin (Cat. No. H107) were purchased from Sigma Aldrich, Millipore Sigma, USA.

### Animals, Ethics, in vivo Efficacy and Durability Studies

All animal procedures complied with AAALAC guidelines and were approved by UCLA institutional animal care and use committees (IACUC) under protocol ARC-2019-011. Mice were housed under controlled temperature and 12-h light/dark cycles with ad libitum access to water and diet. All islets were obtained from deceased donors following consent from next of kin.

Only male hemizygous h-TG mice and age-matched FvB wild-type (WT) mice were used for in vivo efficacy studies because female h-TG mice exhibit estrogen-related protection (*66*). All animals were maintained on a high-fat diet (HFD) to induce obesity and, in h-TG mice, progressive diabetogenic stress. WT mice on HFD served as obese, non-diabetic controls and received sterile PBS (pH 7.4) as vehicle (WT+Vehicle; n = 13–14). h-TG mice subjected to diabetogenic stress (DS) were randomized into the following treatment groups: (i) DS receiving vehicle consisting of 8% (w/v) 2-hydroxypropyl-β-cyclodextrin in sterile PBS (pH 7.4) (DS+Vehicle; n = 8–12); (ii) DS receiving Exendin-4 at 20 µg/kg once daily via intraperitoneal (i.p.) administration for 8 weeks (DS+Ex-4; n = 7–8); (iii) DS receiving the PFKFB3 inhibitor AZ67 at 28 mg/kg i.p. once daily for 4 weeks followed by a 4-week treatment withdrawal (DS+AZ67_G1; n = 9); (iv) DS receiving AZ67 at 28 mg/kg i.p. once daily for 8 consecutive weeks (DS+AZ67_G2; n = 9); and (v) DS receiving a combination of AZ67 (28 mg/kg, i.p.) and Exendin-4 (20 µg/kg, i.p.) once daily for 8 weeks (DS+AZ67+Ex-4; n = 7–9).

For the combination treatment group, AZ67 was administered in the morning and Exendin-4 in the evening, with a minimum interval of 6 hours between administrations to minimize the potential for acute drug–drug interactions. IP-GTT, IP-ITT, blood collection, and collection of pancreata were performed according to the experimental timeline shown in Fig. 2A. The DS+AZ67 group comprised 18 mice through the first 4 weeks of treatment and was then divided into two subgroups, DS+AZ67_G1 and DS+AZ67_G2 (n = 9 each). Data for the comparator analysis were obtained after 4 weeks of treatment and compared across all treatment arms, whereas the durability analysis compared DS+AZ67_G1 (4 weeks on treatment followed by 4 weeks off treatment) with DS+AZ67_G2 (8 weeks of continuous treatment) at the 8-week study endpoint. Body weights and fasting blood glucose levels were assessed weekly.

### Metabolic Testing and Plasma Insulin

For intraperitoneal glucose tolerance tests (IP-GTT), mice were fasted overnight (15 h) and administered glucose (2 g/kg, i.p.), with blood glucose measured at 0, 30, 60, and 120 min. Area under the curve (AUC) was calculated using the trapezoidal rule. Intraperitoneal insulin tolerance tests (IP-ITT) were performed after a 6-h fast using insulin (0.75 U/kg, i.p.), with glucose measured at 0, 20, 40 and 60 min. Plasma insulin was measured at baseline and 30 min post-GTT by ELISA after 4 weeks of treatment. Plasma insulin concentrations were quantified using a commercially available mouse insulin enzyme-linked immunosorbent assay (ELISA) kit (Mercodia Mouse Insulin ELISA; Mercodia AB, Uppsala, Sweden; Catalog No. 10-1247-01), according to the manufacturer’s instructions. Blood samples were collected via the sub-mandibular route, into Sarstedt Microvette CB 300 tubes coated with lithium heparin, at baseline (fasted) and 30 min following intraperitoneal glucose administration during glucose tolerance testing. Samples were centrifuged at 2,000 × g for 10 minutes at 4 °C, and plasma was stored at −80 °C until analysis.

### In Vitro Assays

In vitro assays including ADME studies were performed at Eurofins Discovery. Inhibitory concentrations were determined using the ADP-Glo assay (Promega) as per manufacturer’s protocol, using 60 nM PFKFB3 (produced by Eurofins), 100 μM F6P, and 8 μM ATP in 50mM Trizma Base (pH 8.0), 10mM MgCl_2_, and 5mM sodium phosphate. Binding of inhibitor to PFKFB3 was performed using Spectral Shift. Briefly, PFKFB3 was labelled with RED-NHS (NanoTemper) and incubated (at 15 nM) with test inhibitors in 12-point semi-log dilution series for 1 hour at RT. Spectral Shift was detected on the NanoTemper Dianthus NT23, and Fluorescence ratios (670/650 nm) were calculated using DI.Screening Analysis software (NanoTemper). Dissociation constants (KD) were obtained by fitting dose–response curves to a 1:1 binding model based on the law of mass action (*43*).

### Plasma Protein Binding

Plasma protein binding of AZ67 and AZ26 was determined using an equilibrium dialysis method in a 96-well format (Eurofins Discovery). The compound (10 µM, 1% DMSO final concentration) was incubated with plasma protein matrices in one compartment of a Teflon dialysis block, while the opposing compartment contained phosphate-buffered saline (PBS, pH 7.4). Equal volumes of protein matrix and buffer were loaded, and plates were incubated at 37 °C for 4 h to achieve equilibrium. Following incubation, samples from both compartments were collected, diluted with PBS, precipitated with acetonitrile, and centrifuged. Supernatants were analyzed by HPLC–MS/MS using selected reaction monitoring. Control samples (without dialysis) were processed in parallel to assess recovery. Reference compounds representing low, medium, and high plasma protein binding (acebutolol, quinidine, and warfarin) were included in each assay. Protein binding (%) was calculated from analyte peak areas in protein and buffer compartments, and recovery (%) was determined to assess assay reliability.

### Blood and Plasma Stability Assay

The stability of AZ67 and AZ26 in blood or plasma was assessed using a time-course assay in a 96-well plate format. Test compounds were incubated at 1 µM (0.5% DMSO final concentration) in pre-warmed blood or plasma at 37 °C. Samples were collected at 0, 0.5, 1, 1.5, and 2 h and quenched by transfer into acetonitrile. Following protein precipitation and centrifugation, supernatants were analyzed by HPLC–MS/MS. Compound stability was expressed as the percentage remaining relative to time zero. The apparent half-life (t_1/2_) was determined from the slope of the log-transformed compound concentration versus time, assuming first-order kinetics. Reference compounds (propoxycaine and propantheline) were included to validate assay performance (*67*).

### Human Islet Microtissue Studies

Human islet microtissues experiments were conducted by InSphero using 3D human islet microtissues generated from cryopreserved islets isolated from a male donor (33 years; HbA1c 5.7%). Human islets were purchased from Prodo Laboratories Inc. (Irvine, CA). Dispersed islets were reaggregated into microtissues using the Akura™ PLUS Spheroid Hanging Drop System for 5 days (InSphero AG, CS-06-001-02), to achieve a volume of roughly 1 islet equivalent (IEQ) per MT. The reaggregated islets were transferred to and cultured in Akura™ 96 Spheroid Microplates (InSphero AG, CS-09-001-03) with 3D InSight™ Human Islet Maintenance Medium (InSphero AG, CS-07-005-01). To model disease, human islet microtissues were chronically exposed to glucotoxic stress (11 mM glucose) or cytokine stress (IL-1β, IFNγ, and TNFα) for 6 days. AZ67 and AZ26 were tested at 1 µM and 5 µM. Glucose-stimulated insulin secretion (GSIS) was measured under basal (2.8 mM) and stimulated (16.7 mM) glucose conditions for 120 min. Insulin and proinsulin were quantified by ELISA, and proinsulin:insulin ratios were calculated on a molar basis. hIsMT were fixed in 4% paraformaldehyde, permeabilized, and immunostained for insulin, glucagon, and PFKFB3. The primary antibodies were incubated overnight at 4°C and species-appropriate secondary antibodies were applied for 4 h at room temperature. The following primary antibodies and secondary antibodies were used: rabbit anti-PFKFB3 (Abcam ab181861, Cambridge, MA, USA, 1:100), guinea pig anti-insulin (Abcam, ab195956; 1:100), mouse anti-glucagon (Sigma-Aldrich, G2654; 1:500), F(ab′)₂ fragment donkey anti–guinea pig IgG (H+L) conjugated to fluorescein isothiocyanate (FITC) (Jackson ImmunoResearch, 706-096-148; 1:200), F(ab’)2 conjugates with Alexa 647 donkey anti-mouse IgG (H + L) (Jackson ImmunoResearch 715-606-150, West Grove, PA, USA, 1:200) and F(ab′)₂ fragment donkey anti–rabbit IgG (H+L) conjugated to Cy3 (Jackson ImmunoResearch, 711-165-151; 1:200). Z-stack images were acquired by confocal microscopy using 20x objective with 0.5 µm z-step. During imaging the green (insulin) and far-red channel (glucagon) were set on auto-exposure to capture the insulin and glucagon positive cells from the core of organoid while the orange/yellow channel (PFKFB3) was set at a pre-defined exposure time decided based on signal from healthy control hIsMT. Images were analyzed using QuPath for automated quantification of representative optical z-sections sampled from the top, middle, and bottom regions of each hIsMT, using a validated thresholding pipeline with a fixed threshold range across treatment groups and manual visual inspection. Quantified populations included PFKFB3^+^ INS^+^ β-cells, PFKFB3^+^ GCG^+^ α-cells, and PFKFB3^+^ INS⁻/GCG⁻ stressed endocrine cells. Images from the middle optical z-section of representative hIsMT, encompassing an expanded cross-sectional area, are shown in Fig. 5A,B, whereas the corresponding top and bottom optical z-sections are shown in Figs. S8 and S9. One-way ANOVA followed by Dunnett’s multiple-comparisons test was used for prespecified comparisons of each treatment group with the corresponding stress control. Exploratory pairwise comparisons were additionally performed using unpaired two-tailed t-tests. Data are presented as mean ± SEM, with statistical significance defined as P < 0.05.

### Next-Generation Sequencing and Analysis

At study completion, human islet hIsMT were washed in PBS and lysed in RLT buffer (Qiagen) containing 1% β-mercaptoethanol. Total RNA was isolated using the RNeasy Plus Micro Kit (Qiagen), and samples with RIN >7 were used for analysis. Libraries were prepared using the SMART-Seq HT Kit (Takara Bio) and sequenced (2 x 150 bp) on an Illumina NovaSeq platform, achieving >10 million mapped reads per sample.

Reads were trimmed with Trimmomatic and aligned to the human genome using STAR, and gene counts were generated with featureCounts. Quality control was performed using FastQC, and samples with <10 million mapped reads or <50% mapping efficiency were excluded. The counts were normalized using DESeq2. Differential expression analysis was performed using a negative binomial model with Benjamini-Hochberg correction (FDR <0.05), with |log₂ fold change| >1.5 considered strongly regulated.

Principal component analysis and hierarchical clustering were conducted on the top 2,000 most variable genes. Gene set enrichment analysis (GSEA) was performed using pre-ranked log₂ fold changes with MSigDB gene sets, with FDR <0.1 considered significant. Analyses were conducted in R using Bioconductor packages.

### Pancreatic Histology and Immunofluorescence

Pancreatic tissues were harvested at study termination, fixed in 4% paraformaldehyde overnight at 4°C, processed, and embedded in paraffin by the Translational Pathology Core Laboratory at UCLA. Serial sections (4 µm) for further analysis were prepared across the entire pancreas to enable quantification of injured and dysfunctional β-cell populations, as previously described (*29*). Immunofluorescence imaging was performed on a Zeiss Axio Imager 2.0 microscope equipped with an Apotome structured illumination module and images analysed using QuPath software. Rabbit anti-PFKFB3 (Abcam ab181861, Cambridge, MA, USA, 1:100), Mouse anti-MAIP1 (D-12, 1:50; Santa Cruz Biotechnology) and Guinea pig anti-insulin (Abcam, ab195956; 1:400) were used as primary antibodies. Secondary antibodies were applied for 1h at room temperature and included F(ab′)₂ fragment donkey anti–rabbit IgG (H+L) conjugated to Cy3 (Jackson ImmunoResearch, 711-165-151; 1:200), F(ab’)2 conjugates with Alexa 647 donkey anti-mouse IgG (H + L) (Jackson ImmunoResearch 715-606-150, West Grove, PA, USA, 1:200) and F(ab′)₂ fragment donkey anti–guinea pig IgG (H+L) conjugated to fluorescein isothiocyanate (FITC) (Jackson ImmunoResearch, 706-096-148; 1:200). Nuclei were counterstained and slides were mounted using VECTASHIELD antifade mounting medium containing DAPI (Vector Laboratories, H-1200).

### STATISTICAL ANALYSIS

Statistical analyses followed CRO SOPs and standard biostatistical practices. One-way ANOVA followed by Dunnett’s multiple-comparisons test was used for prespecified comparisons of each treatment group with the corresponding stress control. Exploratory pairwise comparisons were additionally performed using unpaired two-tailed t-tests. Outliers in hIsMT datasets were excluded using the ROUT method (Q = 5%). Data are presented as mean ± SEM, with statistical significance defined as P < 0.05.

## ACKNOWLEDGMENTS

This work was supported by Metanoia Bio, Inc. (funding ST, KR, and GB), including studies conducted at UCLA under Sponsored Research Agreement ISR No. 2021-0206, ADME and kinome profiling by Eurofins DiscoveryOne, and human islet microtissue studies by InSphero AG the latter supported in part by a Research Collaboration Agreement between Metanoia Bio, Inc. and Novo Nordisk A/S. ST, KR, GB, OG, and AvW were also supported by the National Institute of Diabetes and Digestive and Kidney Diseases (NIDDK) of the NIH under Award Number 1R41DK139901-01, awarded to Metanoia Bio, Inc. with consortium funding to The Regents of the University of California, Los Angeles; NIDDK had no role in study design, data collection, analysis, interpretation, or the decision to publish. The content is solely the responsibility of the authors and does not necessarily represent the official views of the NIH.

We thank Dr. Tatyana Gurlo and Dr. Marjan Slak Rupnik for the critical reading of the manuscript.

## Author contributions

- K Raval: Formal analysis of all in vivo and in vitro experiments including immunostaining and imaging, data curation, validation, visualization, and writing, review & editing.
- E Ilegems: Formal analysis of human hIsMT (experimental design) and data interpretation and curation, manuscript draft review and editing.
- A Title: Investigation of human hIsMT, methodology and analysis
- E Fishburn: Formal analysis of the physicochemical properties of AZ67 and AZ26.
- G Bhardwaj: Data curation, software, formal analysis, review and editing.
- O Mirguet: Formal analysis of physicochemical properties of AZ67 and AZ26.
- P Ratcliffe: Supervision of the formal analysis of physicochemical properties of AZ67 and AZ26.
- O Greeff: Formal analysis, and review.
- A van Wyk: Review & editing.
- S Tudzarova: Conceptualization, methodology; validation; formal analysis; investigation; resources; data curation; writing – original draft; writing – review & editing; visualization; supervision; project administration; funding acquisition.

## Competing interests

ST, AvW and OG are scientific co-founders and shareholders of Metanoia Bio, Inc. EF, OM, and PR are employees of Eurofins DiscoveryOne, which performed contract research for Metanoia Bio, Inc. EI is an employee of Novo Nordisk A/S. AT is an employee of InSphero AG, which performed contract research for Metanoia Bio, Inc. The Regents of the University of California hold an equity interest in Metanoia Bio, Inc. as a result of the licensing of intellectual property to the Company. The PFKFB3 inhibitors AZ67 and AZ26 are the subject of patent applications by Metanoia Bio, Inc. and The Regents of the University of California. All other authors declare no competing interests.

## Data and materials availability

All data needed to evaluate the conclusions are present in the paper or Supplementary Materials. All raw data, analytical pipelines, and processed datasets are available from UCLA and Metanoia Bio upon reasonable request, subject to confidentiality and data-sharing agreements with Eurofins, InSphero, and Novo Nordisk A/S.

## SUPPLEMENTARY FIGURES AND FIGURE LEGENDS

**Fig. S1.**
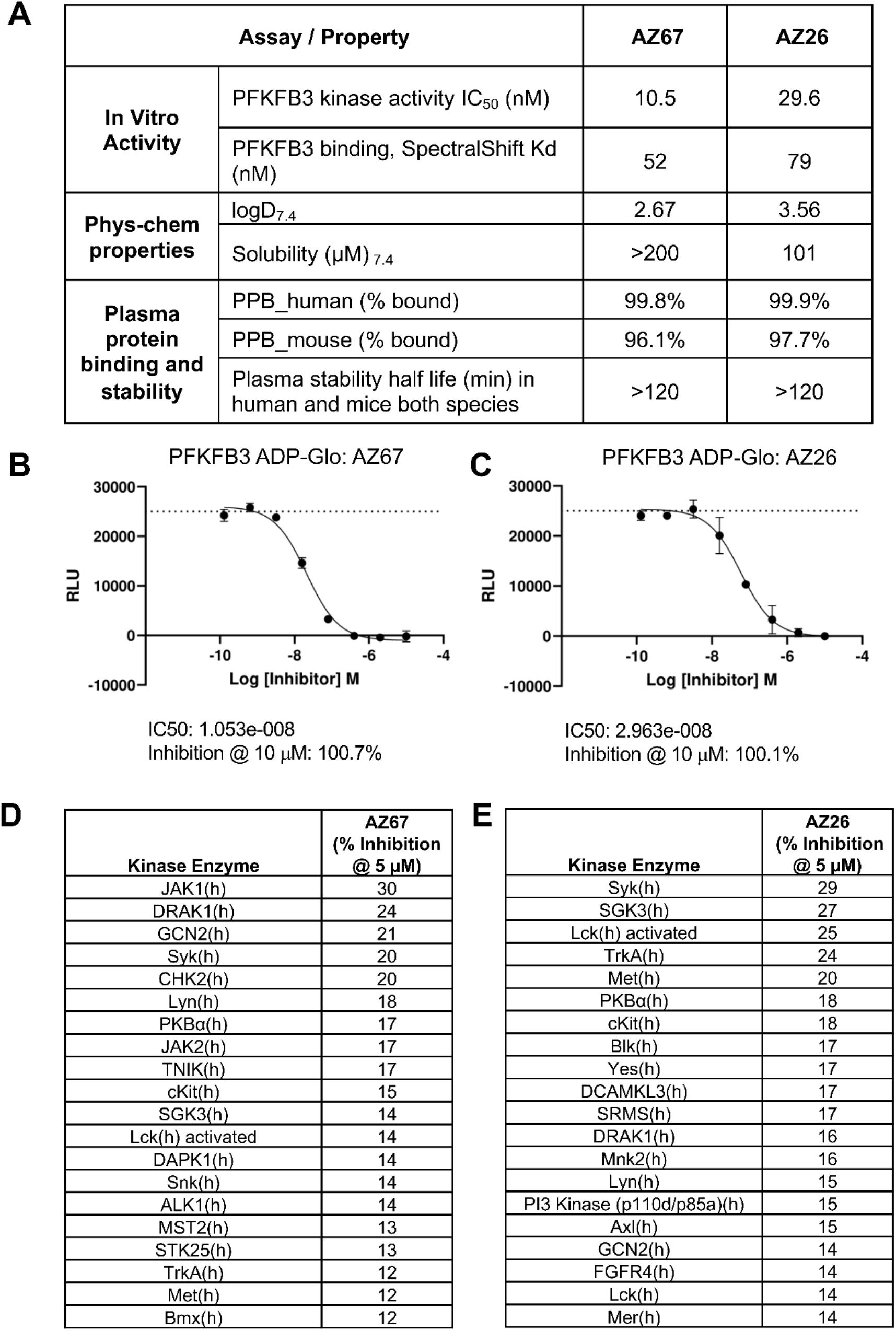
ADME characterization and kinase selectivity profile of AZ67 and AZ26. **(A)** Table summarizing in vitro PFKFB3 inhibitory activity (IC_50_), physicochemical properties, human and mouse plasma protein binding (PPB h and PPB m, respectively), and plasma stability profiles of AZ67 and AZ26 in human and mouse plasma. **(B)** Eight-point ADP-Glo dose-response curve and corresponding IC_50_ values for PFKFB3 inhibition by AZ67. **(C)** Eight-point ADP-Glo dose-response curve and corresponding IC_50_ values for PFKFB3 inhibition by AZ26. **(D)** Kinome profiling analysis showing the top 20 kinases inhibited by AZ67 at 5 μM concentration. **(E)** Kinome profiling analysis showing the top 20 kinases inhibited by AZ26 at 5 μM concentration. Both AZ67 and AZ26 demonstrated high affinity for PFKFB3 with limited off-target kinase activity.

**Fig. S2.**
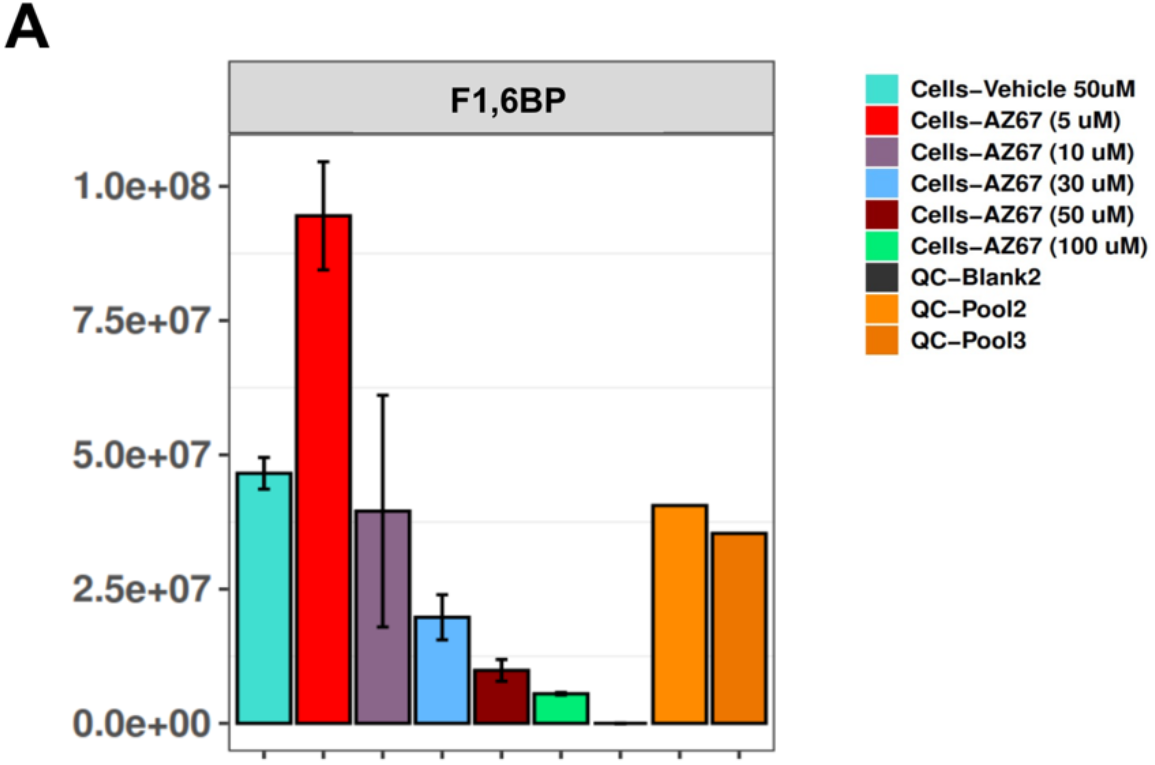
Cellular pharmacodynamic activity of AZ67. Dose-dependent inhibition of PFKFB3 activity by AZ67 in MiaPaCa cells measured by LC-MS/MS quantification of fructose-1,6-bisphosphate (F1,6BP) as a downstream metabolic readout of PFKFB3 pathway activity.

**Fig. S3.**
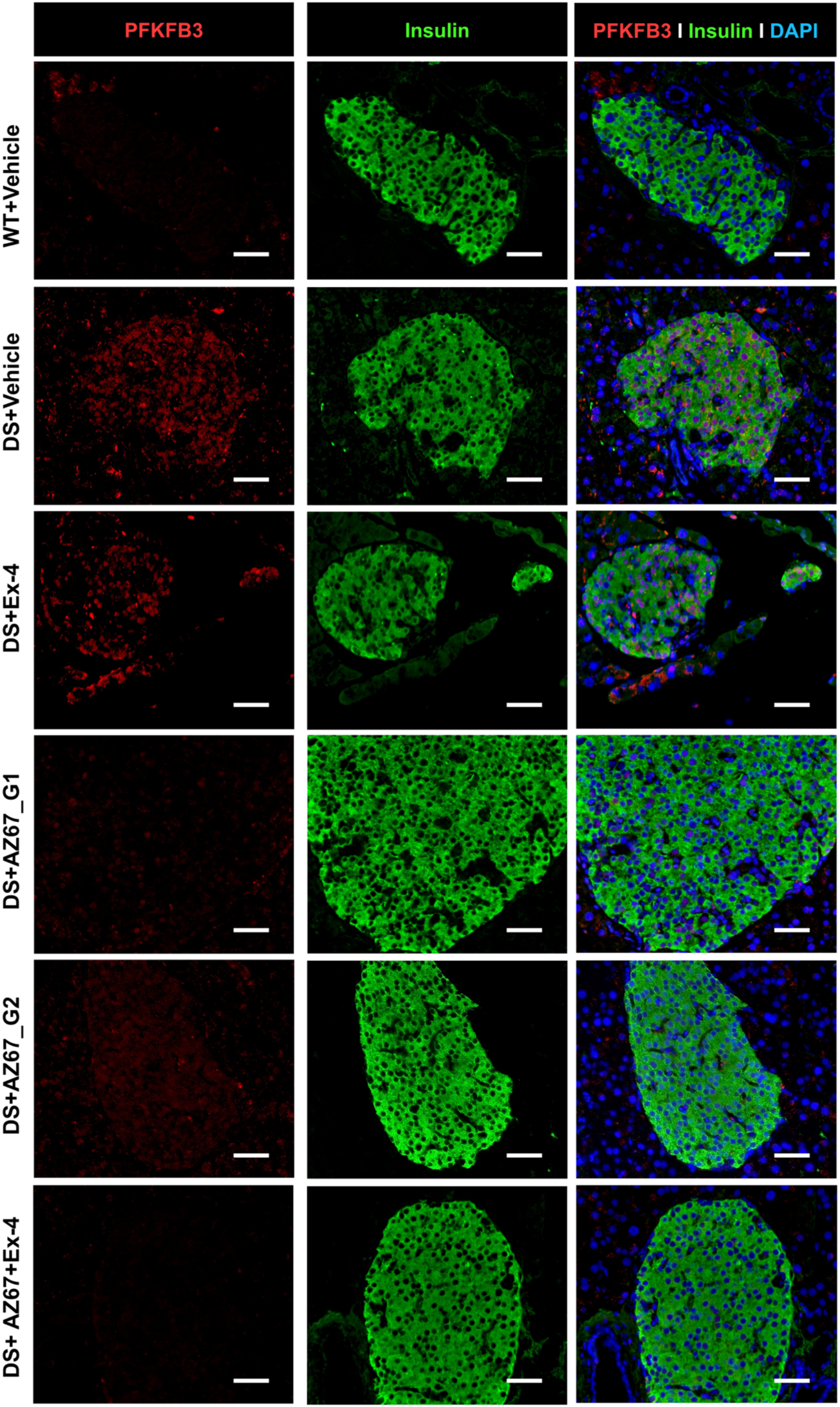
PFKFB3 immunostaining of pancreata from h-TG+HFD mice under different treatments presented in Fig. 3A. Images of split channels from merged images shown in Fig. 3A. Scale bar: 50 μm.

**Fig. S4.**
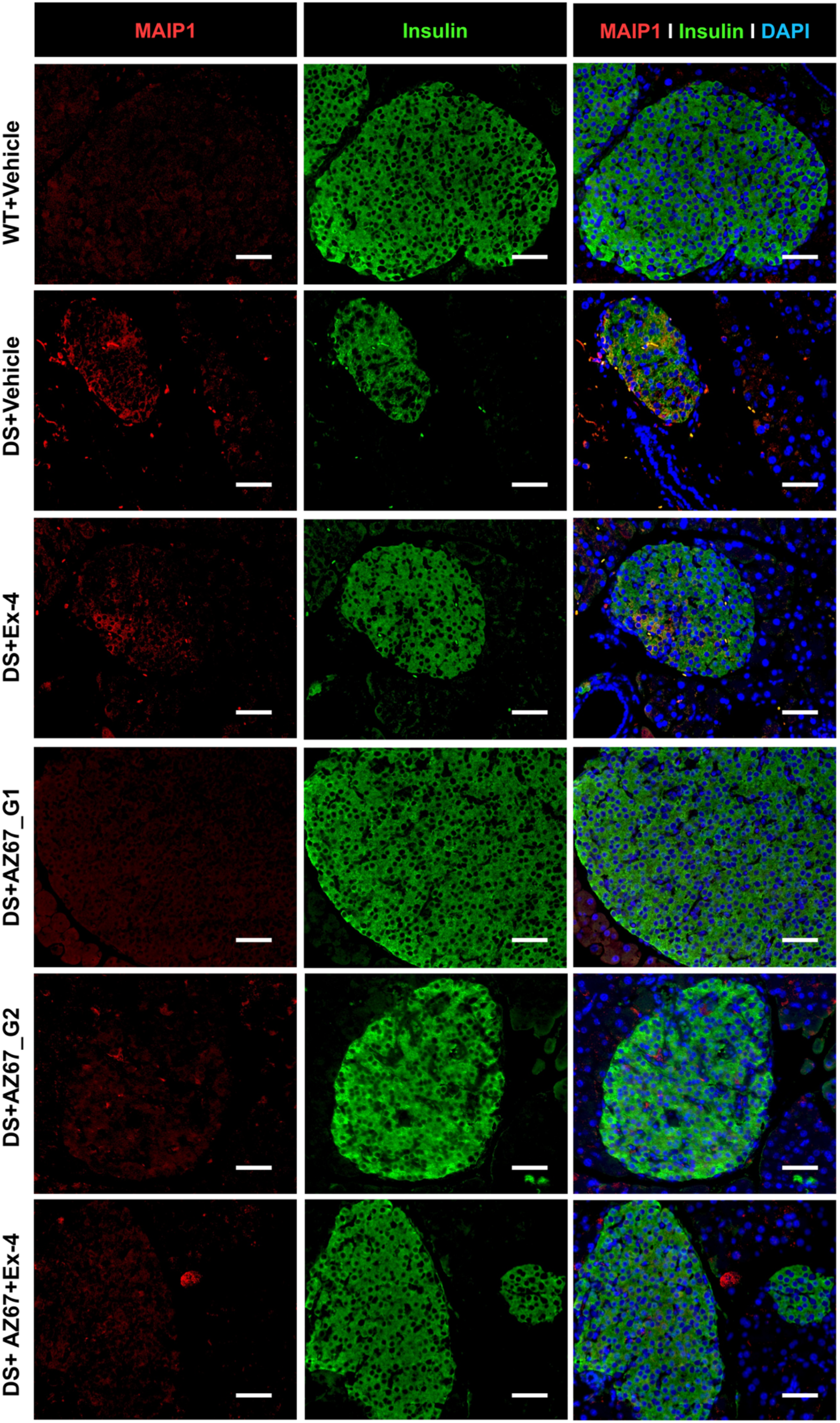
MAIP1 immunostaining of pancreata from h-TG+HFD mice under different treatments presented in Fig. 3B. Images of split channels from merged images shown in Fig. 3B. Scale bar: 50 μm.

**Fig. S5.**
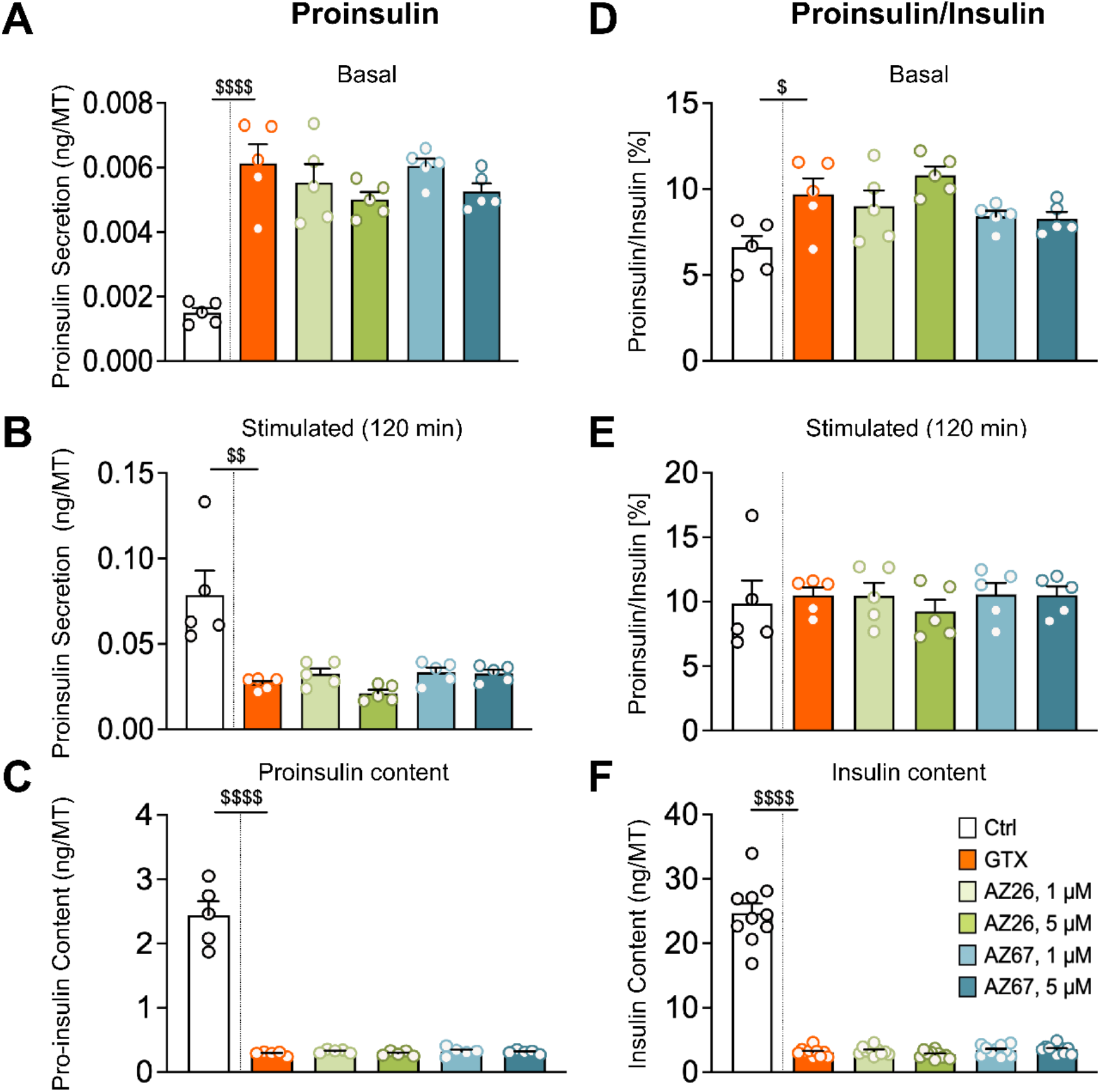
AZ67 and AZ26 do not significantly alter proinsulin processing under glucotoxic (GTX) stress. Human islet microtissues (hIsMTs) were cultured under glucotoxic conditions (GTX, 11 mM glucose) for 6 days in the presence or absence of AZ67 or AZ26 (1 μM or 5 μM). **(A)** Basal proinsulin secretion per hIsMT. **(B)** Glucose-stimulated proinsulin secretion per hIsMT following 120-minute stimulation with 16.7 mM glucose. **(C)** Total proinsulin content normalized to microtissue. **(D)** Basal proinsulin-to-insulin ratio per hIsMT. **(E)** Stimulated proinsulin-to-insulin ratio per hIsMT following 120-minute glucose stimulation. **(F)** Total insulin content normalized to microtissue. Data are presented as mean ± SEM (n = 5–10 independent microtissues per group). Ratios were calculated using molar concentrations. Following outlier removal (ROUT analysis, Q = 5%), one-way ANOVA followed by Dunnett’s multiple-comparisons test was performed relative to the GTX stress group (stars). Direct pairwise comparisons between GTX and individual treatments were additionally assessed using unpaired two-tailed t-tests (hash symbols). Unpaired t-tests comparing healthy CTRL and GTX microtissues were used to validate stress induction (dollar symbols). Healthy control microtissues were cultured under standard glucose conditions (2.8 mM) versus glucotoxic conditions (11 mM). P < 0.05 (*/#/$), P < 0.01 (**/##/$$), P < 0.001 (***/###/$$$), and P < 0.0001 (****/####/$$$$).

**Fig. S6.**
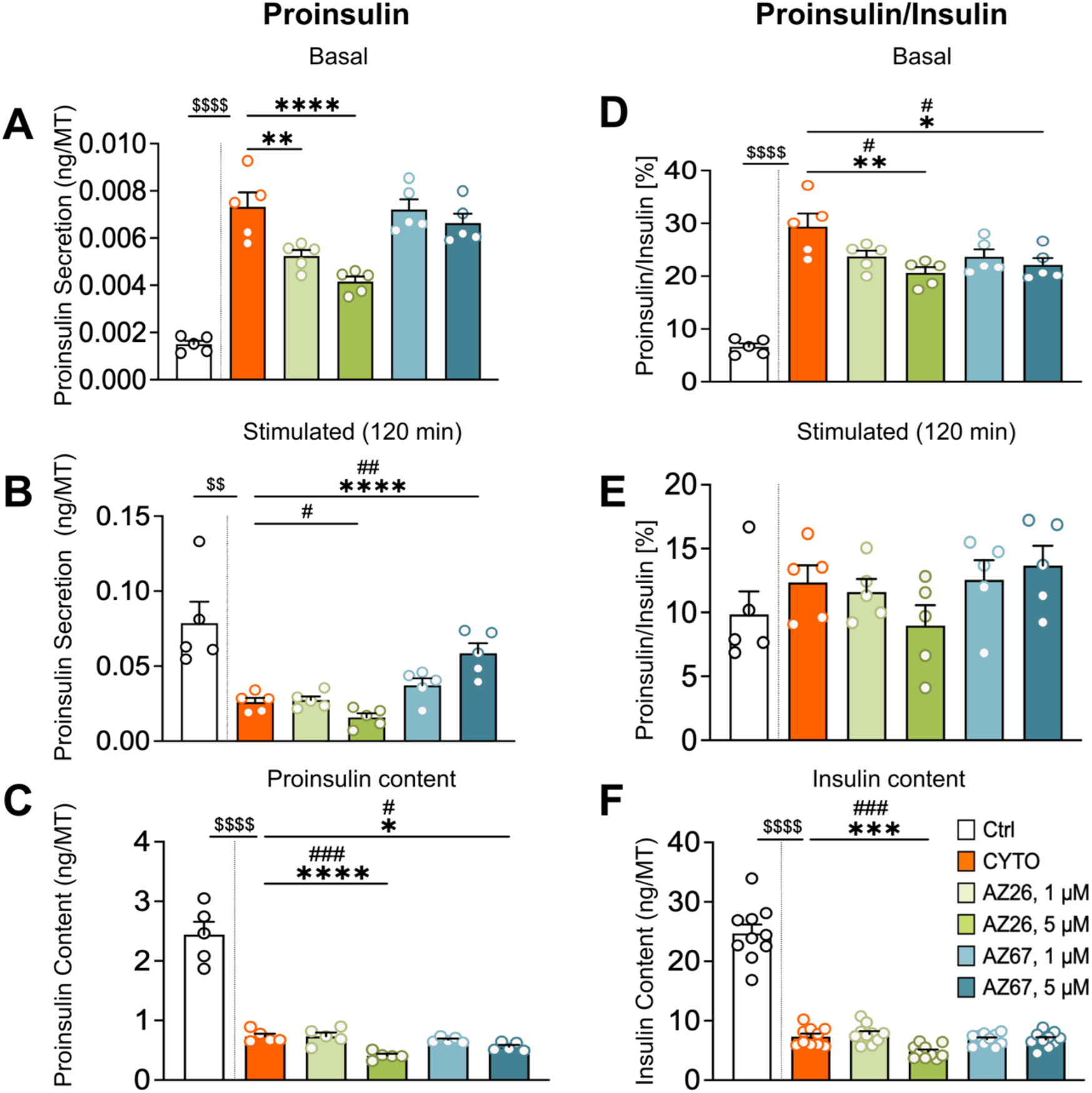
AZ67 increases basal proinsulin processing in cytokine (CYTO) stress. Human islet microtissues were cultured for 6 days under cytokine-induced stress (2 ng/mL IL-1β, 10 ng/mL TNFα, and 10 ng/mL IFNγ) in the presence or absence of AZ67 or AZ26 (1 or 5 μM). **(A)** Basal proinsulin secretion per hIsMT. **(B)** Glucose-stimulated proinsulin secretion per hIsMT following 120-minute stimulation with 16.7 mM glucose. **(C)** Total proinsulin content normalized to microtissue. **(D)** Basal proinsulin-to-insulin ratio per hIsMT. **(E)** Stimulated proinsulin-to-insulin ratio per hIsMT following 120-minute glucose stimulation. **(F)** Total insulin content normalized to microtissue. Data are presented as mean ± SEM (n = 5–10 independent microtissues per group). Ratios were calculated using molar concentrations. Following outlier removal (ROUT analysis, Q = 5%), one-way ANOVA followed by Dunnett’s multiple-comparisons test was performed relative to the CYTO stress group (stars). Direct pairwise comparisons between CYTO and individual treatments were additionally assessed using unpaired two-tailed t-tests (hash symbols). Unpaired t-tests comparing healthy CTRL and CYTO microtissues were used to validate stress induction (dollar symbols). P < 0.05 (*/#/$), P < 0.01 (**/##/$$), P < 0.001 (***/###/$$$), and P < 0.0001 (****/####/$$$$).

**Fig. S7.**
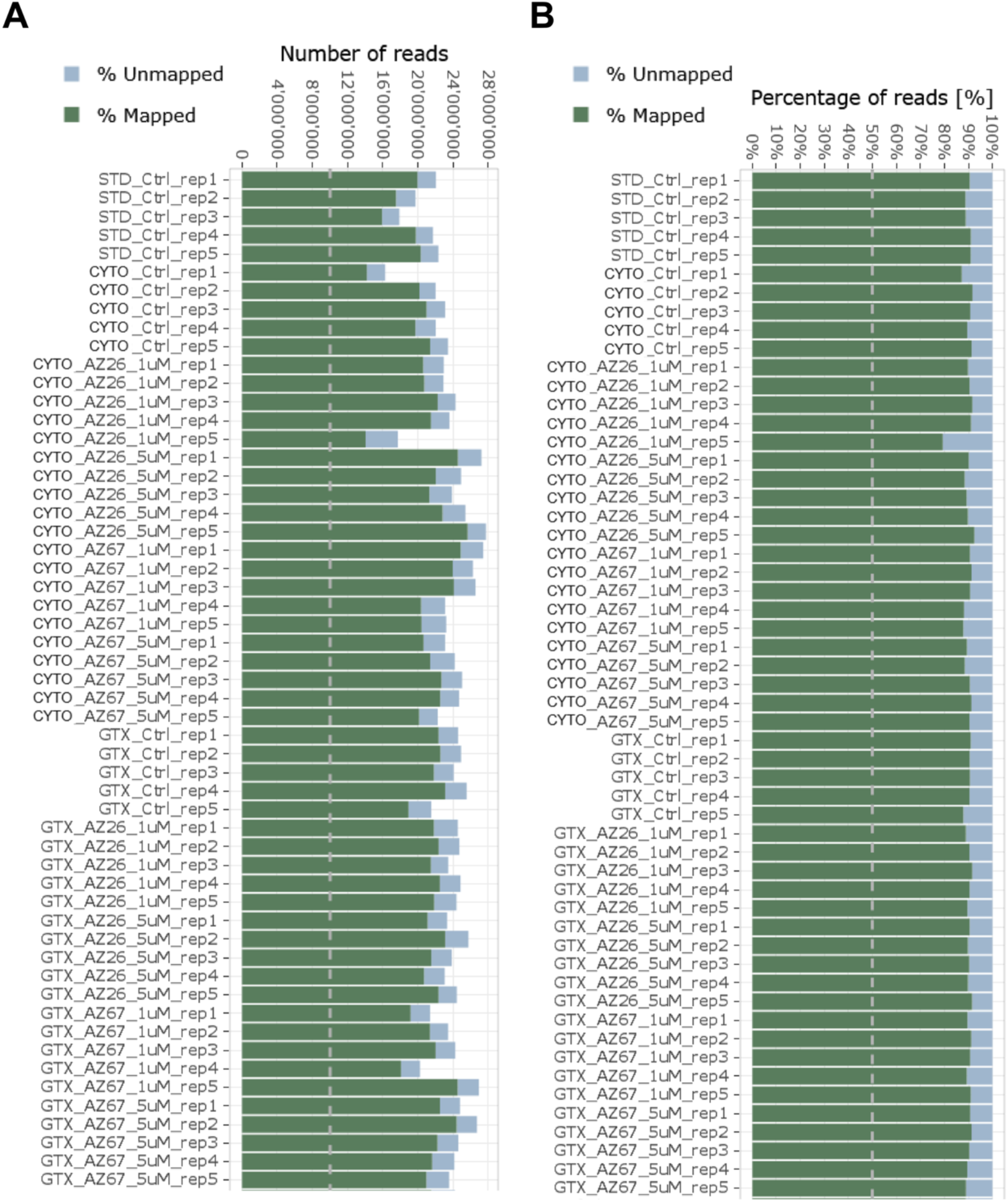
Human islet microtissues’ RNA-Seq quality control. **(A)** Percentage of mapped and unmapped sequencing reads across all samples and treatment groups. Mapping rates were high and consistent across conditions, with an average of ∼90%, indicating robust alignment quality. **(B)** Total sequencing depth per sample, showing the number of mapped and unmapped reads. Average sequencing depth was ∼23.5 million reads per sample, exceeding the predefined threshold and demonstrating consistent library complexity across conditions. All samples passed quality control criteria (>10 million mapped reads and >50% mapping efficiency) and were included in downstream analyses.

**Fig. S8.**
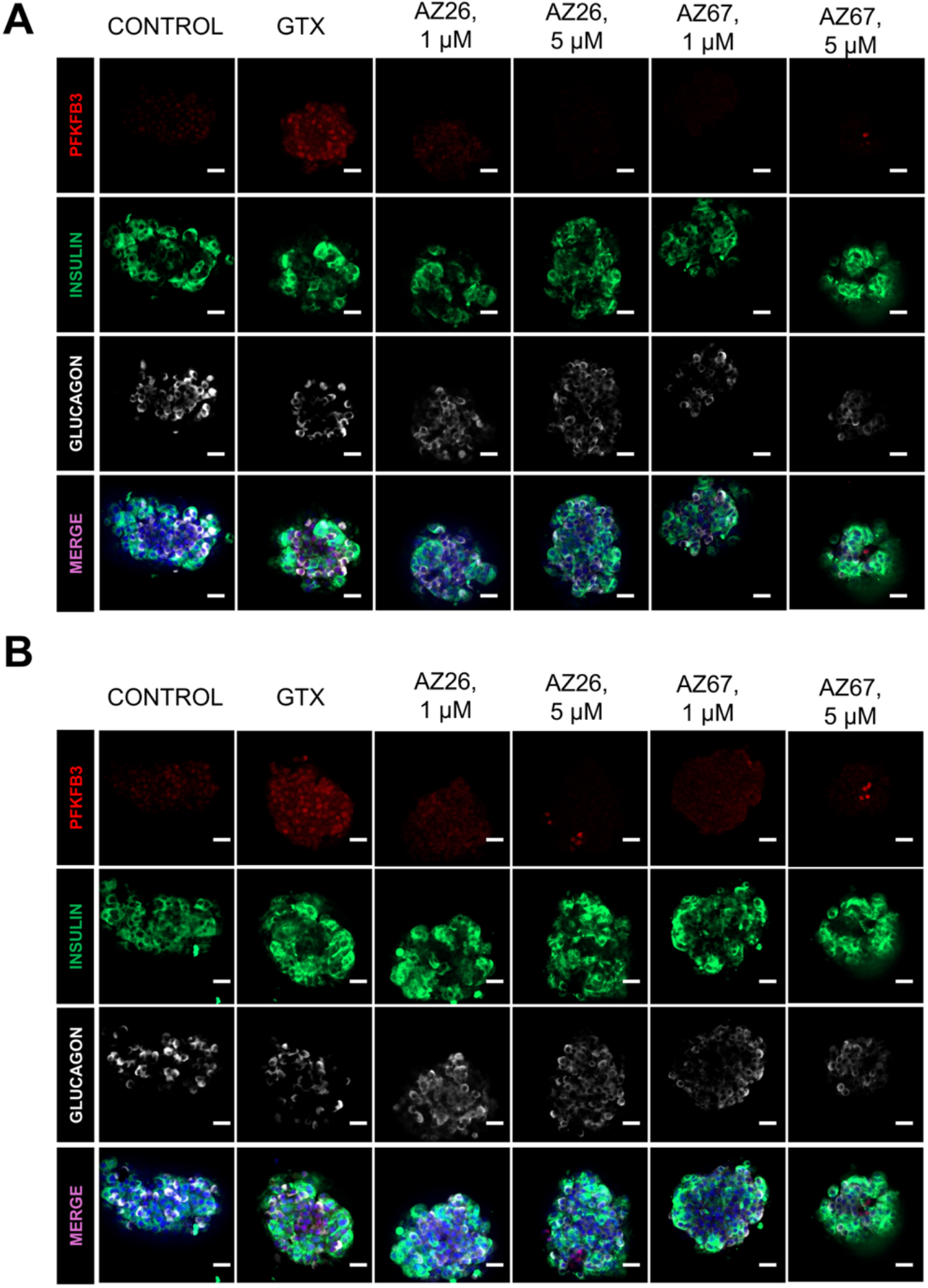
Representative immunofluorescence images of top and bottom optical z-sections of human islet microtissues from the glucotoxicity group shown in Fig. 5A. Human islet microtissues (hIsMT) were stained for insulin (green), glucagon (white), PFKFB3 (red), and nuclei (blue). Representative optical z-sections from (A) the top and (B) the bottom regions of the hIsMT are shown. The corresponding middle optical z-sections are presented in Fig. 5A. Scale bar, 20 μm.

**Fig. S9.**
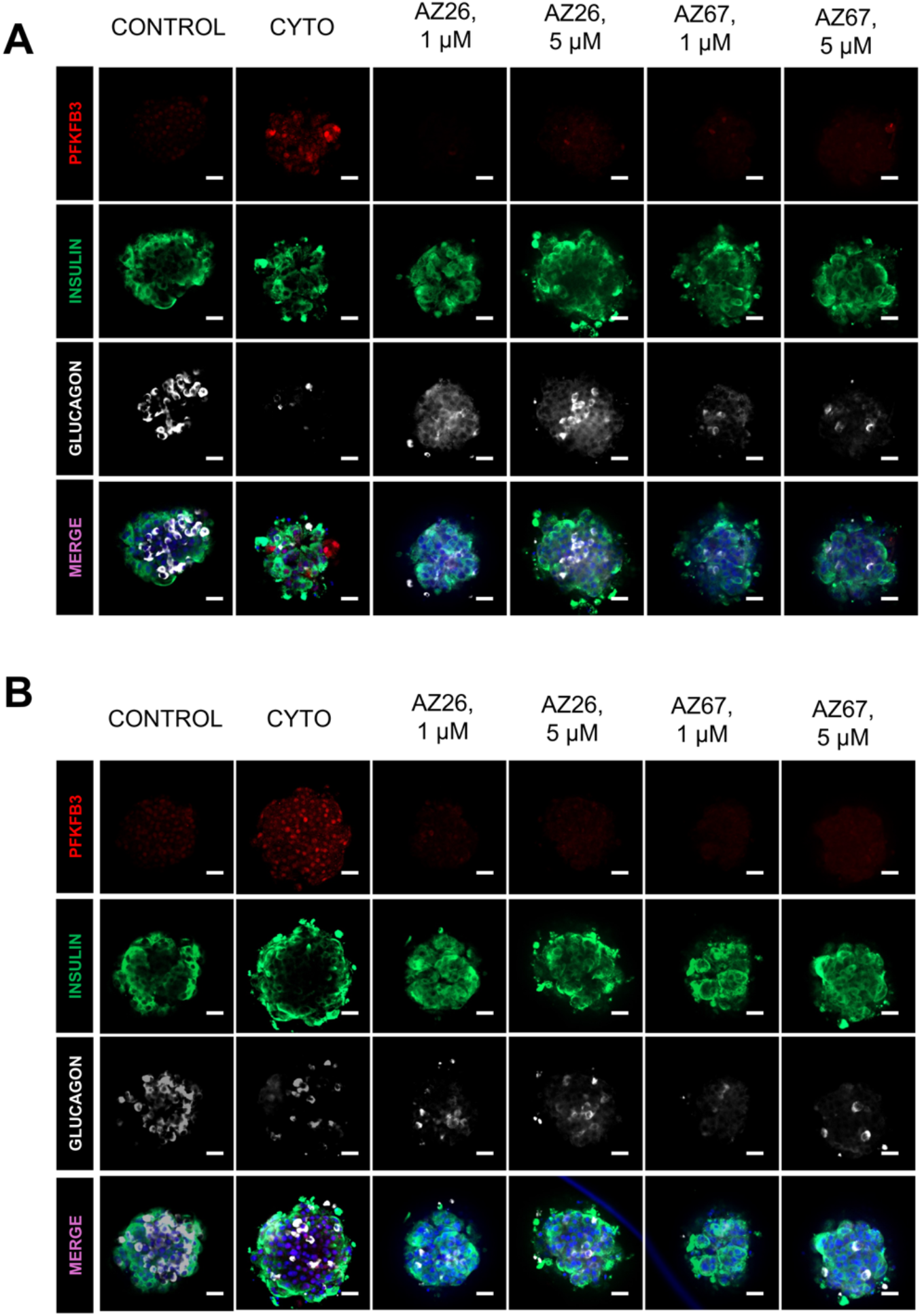
Representative immunofluorescence images of top and bottom optical z-sections of human islet microtissues from the cytokine-stress group shown in Fig. 5B. Human islet organoid microtissues (hIsMT) were stained for insulin (green), glucagon (white), PFKFB3 (red), and nuclei (blue). Representative optical z-sections from (A) the top and (B) the bottom regions of the hIsMT are shown. The corresponding middle optical z-sections are presented in Fig. 5B. Scale bar, 20 μm.

**Fig. S10.**
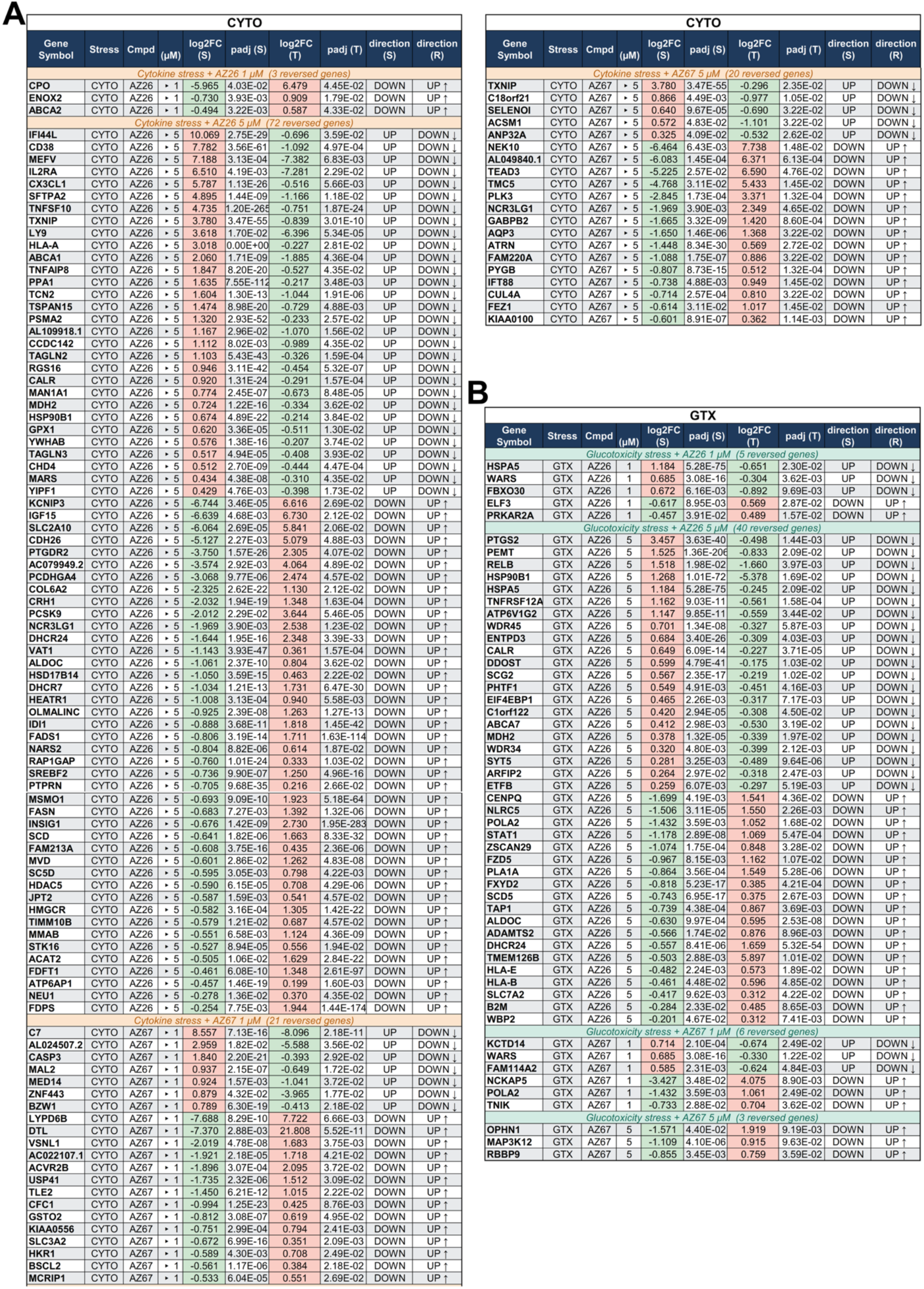
Reversed genes in (A) CYTO-stressed and (B) GTX-stressed human islet hIsMT after AZ67 and AZ26 treatment. DEGs reversed by AZ26 or AZ67 in CYTO and GTX-stressed (S) hIsMT, respectively. Reversal (rescue, R) requires significance (Padj ≤ 0.05) in both stress and treatment (T) comparisons with opposite log2FC sign. HIsMT were exposed 6 days to GTX (11 mM glucose) or CYTO (IL-1β, TNFα, IFNγ) then treated with AZ26 or AZ67 (1 or 5 µM). Green/red arrows indicate rescued/suppressed genes. A total of 170 gene-reversal events, corresponding to 160 unique genes, were identified across all conditions.

